# TLR4-dependent signaling drives extracellular catabolism of low-density lipoprotein aggregates

**DOI:** 10.1101/610162

**Authors:** Rajesh K. Singh, Abigail S. Haka, Arky Asmal, Valéria C. Barbosa-Lorenzi, Inna Grosheva, Harvey F. Chin, Yuquan Xiong, Timothy Hla, Frederick R. Maxfield

## Abstract

**Objective:** Aggregation and modification of low-density lipoproteins (LDL) promotes their retention and accumulation in the arteries. This is a critical initiating factor during atherosclerosis. Macrophage catabolism of aggregated LDL (agLDL) occurs using a specialized extracellular, hydrolytic compartment, the lysosomal synapse (LS). Compartment formation by local actin polymerization and delivery of lysosomal contents by exocytosis promotes acidification of the compartment and degradation of agLDL. Internalization of metabolites such as cholesterol promotes foam cell formation, a process that drives atherogenesis. Further, there is accumulating evidence for the involvement of TLR4 and its adaptor protein MyD88 in atherosclerosis. Here, we investigated the role of TLR4 in catabolism of agLDL using the LS and foam cell formation.

**Approach and Results:** Using bone marrow-derived macrophages (BMMs) from knockout mice, we find that TLR4 and MyD88 regulate compartment formation, lysosome exocytosis, acidification of the compartment and foam cell formation. Using siRNA, pharmacological inhibition and knockout BMMs, we implicate SYK, PI3 kinase and Akt in agLDL catabolism using the LS. Using bone marrow transplantation of LDL receptor knockout mice with TLR4KO bone marrow, we show that deficiency of TLR4 protects macrophages from lipid accumulation during atherosclerosis. Finally, we demonstrate that macrophages *in vivo* form an extracellular compartment and exocytose lysosome contents similar to that observed *in vitro* for degradation of agLDL.

**Conclusions:** We present a mechanism in which interaction of macrophages with agLDL initiates a TLR4 signaling pathway, resulting in formation of the LS, catabolism of agLDL and lipid accumulation *in vitro* and *in vivo*.

## INTRODUCTION

Excessive cholesteryl ester and lipid accumulation in macrophages leading to foam cell formation is a key process during the development of atherosclerosis ^1^. Although mechanisms of cholesterol accumulation have been studied intensively, the *in vivo* contributions of different pathways leading to foam cell formation are not understood. One proposed mechanism of cholesterol delivery is the uptake of modified monomeric low-density lipoprotein (LDL) via various scavenger or pattern recognition receptors ^2, 3^. However, disparate results have been reported using mouse models of atherosclerosis lacking scavenger receptor A1 (SCARA1) and CD36, which are involved in the uptake of oxidized LDL ^4, 5^.

In atherosclerotic plaques, the vast majority of the LDL that interacts with immune cells is not monomeric but rather aggregated and avidly bound to the subendothelial matrix ^6–8^. Mouse models in which the subendothelial retention of LDL is blocked develop significantly less atherosclerosis than their wild-type (WT) counterparts, with lesion area of the total aorta reduced by as much as 63% ^9^. Further, in fatty streaks of the human arteries, over 90% of lesional lipoproteins could not be released by extraction or electrophoresis, indicating their tight association with the vascular wall constituents ^8^. Thus, LDL aggregation and retention is a critical event in atherogenesis and merits consideration in the mechanisms regulating foam cell formation.

Importantly, these aggregated/retained lipoprotein deposits require different cellular uptake processes than those used for endocytosis of monomeric lipoproteins. When macrophages come into contact with LDL aggregates, an extracellular, acidic, hydrolytic compartment (a lysosomal synapse) is formed by a process requiring local actin polymerization ^10–14^. The low pH of the compartment, which confers activity to lysosomal hydrolases delivered by targeted exocytosis, is maintained by V-ATPase in the macrophage plasma membrane ^10^. Following partial catabolism outside the cell, aggregate remnants can be internalized by the macrophage and digested in lysosomes. The lysosomal synapse (LS) allows rapid catabolism of even large aggregates or aggregate complexes that are tightly linked to the extracellular matrix that could not be internalized by phagocytic mechanisms.

In this study, we investigate the receptors and signal transduction pathways involved in the formation and function of the LS. We present a new signaling mechanism for macrophage lipid accumulation in which aggregated LDL (agLDL) is involved in activation of toll-like receptor 4 (TLR4), resulting in local actin polymerization, lysosome exocytosis, and generation of a low pH environment. There is evidence for the involvement of TLR4, a pattern recognition receptor of the innate immune system, in the pathogenesis of atherosclerosis, with both human and animal studies suggesting that TLR4 is proatherogenic ^15–18^. However, most research on the role of TLR4 in atherosclerosis has focused on innate immunity because TLRs represent the major proximal sensory apparatus by which the host detects the presence of foreign pathogens. Herein, we show that, in addition to its role in innate immunity, TLR4 can regulate lipid accumulation during macrophage catabolism of agLDL, and we demonstrate a signaling cascade involving low-density lipoprotein receptor-related protein (LRP-1), the toll-like receptor adaptor molecule myeloid differentiation primary response 88 (MyD88), spleen tyrosine kinase (SYK), CD14, phosphoinositide 3-kinase (PI3K) and Akt. We show that both *in vitro* and *in vivo*, macrophages lacking TLR4 are protected against rapid internalization of free cholesterol and ensuing foam cell formation and provide *in vivo* evidence that macrophages use lysosome exocytosis to the LS for the degradation of agLDL.

## MATERIALS AND METHODS

### Animals

Mice were housed in a pathogen-free environment at Weill Cornell Medical College and used in accordance with protocols approved by the Institutional Animal Care and Utilization Committees. Male and female knockout mice (*Tlr4*^−/−^, *Myd88*^−/−^, *Cd14*^−/−^, *Ldlr*^−/−^, *Cd36*^−/−^, *Scara1*^−/−^ and *Apoe*^−/−^) and appropriate controls were obtained from Jackson Laboratories and were used to prepare BMMs. Legs from *Akt1*^−/−^ and *Akt2*^−/−^ mice were kindly provided by Dr. Morris Birnbaum. For the examination of the lipid content of macrophages in atherosclerotic plaque, 6 *Ldlr*^−/−^ female mice were purchased from Jackson Laboratories.

### Cells and Cell Culture

Mouse macrophage cell lines J774A.1, RAW264.7 and mouse fibroblast cell line L-929 were obtained from the American Type Culture Collection (Manassas, VA). J774 cells stably expressing control and TLR4 targeting shRNA were kindly provided by Dr. Yuri Miller and were described previously ^19^. Cells were maintained in Dulbecco’s Modified Eagle Medium (DMEM) supplemented with 1% (v/v) penicillin/streptomycin, 3.7 g/L sodium bicarbonate and 10% (v/v) heat-inactivated fetal bovine serum (FBS). ShRNA expressing cells were maintained in the same media supplemented with 1 mg/mL G418. BMMs were cultured as described previously ^20^. For all live cell imaging experiments, media was changed to medium 2 (M2) + 0.2% glucose (w/v).

### Reagents

CypHer 5E Mono N-hydroxysuccinimide ester (CypHer 5E) was purchased from GE Healthcare (Chalfont St. Giles, U.K.). AlexaFluor488 (A488), A546, A633, LipidTOX green, A488-phalloidin, biotin-fluorescein-dextran (10,000 MW) and aminophenyl fluorescein (APF) were purchased from Invitrogen (Carlsbad, CA). DNase I was purchased from New England BioLabs (Ipswich, MA). BAY 61-3606 was purchased from EMD Millipore (Darmstadt, Germany). MPLA was purchased from Enzo Life Sciences (Farmingdale, NY). A66, TGX-221, AS-605240 and CAL-101 were purchased from Selleck Chemicals (Houston, TX). All other chemicals were purchased from Sigma Chemicals (St. Louis, MO).

### Lipoproteins

LDL was isolated from fresh human plasma by preparative ultracentrifugation as previously described ^21^. LDL was vortex aggregated for 30 sec to form agLDL. ^22^. AcLDL was prepared by acetylation of LDL with acetic anhydride as described previously ^23^. Levels of endotoxin in the LDL isolated from human plasma, quantified using a chromogenic limulus amebocyte lysate endotoxin assay kit (GenScript, Piscataway, NJ), were significantly below those necessary for activation of TLR4 (typically <0.05 ng/ml).

### Antibodies and Immunoblotting

Macrophages were plated into a 6 well plate (0.5 × 10^6^/well) and cultured overnight. Cells were subsequently treated with 1 mg/mL agLDL, 1 mg/mL monomeric LDL (mLDL) or 100 ng/mL LPS in standard culture medium for various periods of time. Cells were lysed by scraping in 400 μL ice cold lysis buffer RIPA buffer + 1 x complete mini EDTA-free protease inhibitors (Roche Diagnostics Corporation, Indianapolis, IN) + 1 x halt phosphatase inhibitor (Thermo Scientific, Rockford, IL). Total cell lysates were diluted 1:1 in 2× lamelli buffer, boiled for 10 min and subjected to electrophoresis and immunoblotting. Antibodies used for immunoblotting were: Anti-phospho-Akt (T308) (1:500, #2965), anti-phospho-Akt (S473) (1:1000, #4060) and anti-Akt (pan) (1:1000, #4691) and anti-SYK (1:1000, #2712), all from Cell Signaling Technologies (Danvers, MA). As a loading control, anti-GAPDH (1:2500, #9485) (Abcam, Cambridge, MA) was used. Densitometry quantification of protein expression was performed using ImageJ (NIH, Bethesda, MD).

### Immunostaining

For immunostaining of TLR4, RAW264.7 cells were treated with A546-agLDL for 1 hr, and fixed with 3% (w/v) paraformaldehyde/0.5% (w/v) glutaraldehyde for 45 min. Cells were blocked at room temperature using 10% (v/v) goat serum in PBS for 1 hr. Cells were then stained for TLR4 overnight at 4°C using a rat anti-mouse CD284(TLR4)/MD-2 ab (1:300, #562221) or a rat IgG2a, κ isotype control (1:300, #555840), both from BD Biosciences, San Jose, CA). Cells were stained using A488-anti-rat antibody (Invitrogen) at room temperature using a 1:400 dilution. Next, the plasma membrane was stained using A633-Wheat Germ Agglutinin (Invitrogen) using 1:1000 dilution for 10 min at room temperature. All labeling was carried out in PBS containing 3% goat serum. Cells were then analyzed by confocal microscopy.

### SiRNA transfection

J774 cells were silenced using a pool of 4 different Flexitube siRNAs (Qiagen) for SYK with Lipofectamine RNAiMAX reagent as described previously ^20^. After 48 hours knockdown, J774 cells cultured in a 6 well plate were lysed and SYK protein expression assessed by immunoblot. J774 cells cultured on Poly-D-lysine coated glass-coverslip bottom dishes were used to assess lysosome exocytosis and actin polymerization.

### Actin Measurements

To visualize F-actin, macrophages were incubated with A546-agLDL for 1 hr, washed with PBS and fixed for 15 min with 3% PFA. Cells were subsequently washed with PBS, and incubated with 0.02 U/ml of A488-phalloidin in 0.5% (w/v) saponin in PBS for 1 hr at room temperature. Images were acquired using a Zeiss LSM 510 laser scanning confocal microscope with a 40× 0.8 NA objective. Image quantification was performed using MetaMorph software as described previously ^24^.

### Delivery of Lysosomal Contents

Lysosome labeling of macrophages plated on Poly-D-lysine coated glass-coverslip bottom dishes was accomplished via an 18 hr pulse with 1 mg/ml biotin-fluorescein-dextran. Cells were chased for 2 hours in DMEM and subsequently incubated with streptavidin-A546-agLDL for 90 min. Next, cells were incubated with 200 μM biotin for 10 min in order to bind any unoccupied streptavidin sites prior to cell permeabilization. Cells were then fixed with 1% PFA for 15 min, washed, and permeabilized with 1% Triton X-100 for 10 min. Images were acquired with the confocal microscope described above using a 40× 0.8 NA objective. For image quantification, MetaMorph software was used. All images subjected to comparative quantification were acquired on the same day using the same microscope settings. Each experiment was repeated at least three times. For every experiment >30 randomly chosen fields with a total >200 cells per condition were imaged and subjected to quantification. To quantify the amount of lysosome exocytosis, we obtained a single plane for each field at wavelengths appropriate for streptavidin-A546-agLDL (red) and biotin-fluorescein-dextran (green). We determined a threshold in the red channel (agLDL) that would include nearly all of the observable agLDL in the images. We then measured the total fluorescein fluorescence intensity within the thresholded area for each field. We used the same threshold level for each image within an experimental data set. By this procedure the total fluorescein signal intensity within the thresholded regions per field was measured. Data was normalized by the amount of biotin-fluorescein-dextran delivered to lysosomes as determined by confocal imaging of non-permeabilized cells.

### pH Measurements

Macrophages were plated on Poly-D-lysine coated glass-coverslip bottom dishes. The cells were incubated for 30 min with CypHer 5E, a pH sensitive fluorophore, and A488, a pH insensitive fluorophore, dual labeled agLDL. The pH values within each pixel were assessed quantitatively by comparison with ratio images obtained in calibration buffers of varying pH. Live cells were imaged on the confocal microscope described above using a 63× 1.4 NA objective. Cell temperature was maintained at 37°C with a heated stage and objective heater. Data were analyzed with MetaMorph image analysis software. A binary mask was created using the A488 signal intensity and applied to both channels to remove background noise. Images were convolved with a 7×7 pixel Gaussian filter, and ratio images were generated.

### Foam Cell Formation

Macrophages were plated onto poly-D-lysine coated glass-coverslip bottom dishes overnight and incubated with A546-agLDL or AcLDL for 12 hours at 37°C and 5% CO_2_. Cells were fixed with 3% PFA for 20 min and washed with PBS. They were subsequently stained with LipidTOX green (1:1000) for 15 min at room temperature and washed with PBS. Images were acquired with the confocal microscope described above using a 40× 0.8 NA objective. For image quantification, MetaMorph software was used. Images were thresholded to exclude fluorescence signal not associated with lipid droplets prior to quantification of total integrated LipidTOX green signal per field.

### ROS production

To examine the production of ROS in the lysosomal synapse, J774 cells plated on poly-D-lysine coated glass-coverslip bottom dishes were incubated with A546-agLDL for 30 min in DMEM. Media was replaced with M2 + 0.2% glucose (w/v) containing 15 μM APF and live cells were imaged on the confocal microscope described above using a 63× 1.4 NA objective. Cell temperature was maintained at 37°C with a heated stage and objective heater.

### Bone Marrow Transplantation

Bone marrow transplantation was performed by injecting 1×10^7^ total bone marrow cells via tail vein into lethally irradiated (950 rads) 7-week old recipient mice. Bone marrow cells were harvested by flushing femurs and tibias of donor animals including *C57BL/6* and *Tlr4*^−/−^mice. After 5 weeks of bone marrow engraftment, recipient mice were transitioned to a high fat diet (21% milk fat, 0.15% cholesterol; Harlan Teklad 88137) and maintained on the diet for 16 weeks. Mice were then euthanized via carbon dioxide inhalation.

### Lipid analysis of macrophages isolated from murine atherosclerotic plaque

Aortic single cells were prepared as described previously ^13^ and cytospun onto glass slides. After blocking with 10% goat serum for 1 hour, macrophages were identified using a rat monoclonal antibody for F4/80 (Abcam ab6640, Cambridge, MA) at 1:300 dilution overnight at 4°C and AlexaFluor546 (A546) anti-rat secondary antibody (Invitrogen) at 1:400 dilution for 2 hours at room temperature. All antibody labeling was carried out in PBS containing 3% goat serum. Next, samples were stained with 1 ¼g/ml Hoechst for 10 min. at room temperature. Last, all samples were stained with LipidTOX green 1:1000 in PBS for 20 min. Images were acquired with a Zeiss LSM 510 laser scanning confocal microscope (Thornwood, NY) using a 63× 1.4 NA objective.

### Adoptive transfer of biotin-fluorescein-dextran labeled bone marrow derived macrophages to ApoE^−/−^ mice

Apolipoprotein E null (*Apoe*^−/−^) mice were placed on a high fat diet (21% milk fat, 0.15% cholesterol; Harlan Teklad) for 24 weeks. Mice were injected with streptavidin-A633-LDL or A546-LDL (1 mg/mouse). Wild-type BMMs (4 × 10^6^) were loaded overnight with 1 mg/ml biotin-fluorescein-dextran and adoptively transferred to the mice via tail vein injection the following day. 3 days after, mice were euthanized, perfused with PBS and aortas taken for sectioning.

### Aortic fixation and sectioning

For tissue processing, aortas were fixed overnight in 3% PFA at 4°C. Fixed aortas were placed in a solution of 30% sucrose in PBS and stored at 4°C overnight. Aortas were then gently agitated in embedding media (1:2 ratio of 30% sucrose in PBS in optimal cutting temperature (OCT) medium and then frozen in the same media using 2-methylbutane and liquid nitrogen. Samples were then cut into 8 μm sections using a Cryostat, mounted onto glass slides and coverslips attached using Vectorshield mounting medium for fluorescence (Vector Laboratories, Burlingame, CA). For cholesterol visualization, samples were stained with 50 μg/mL of filipin in PBS for 1 hr at room temperature. For F4/80 labeling, sections were stained with primary antibody at 1:500 dilution overnight at 4°C, followed by A546-anti-rat secondary antibody at 1:500 dilution for 4 hours at room temperature. For adoptive transfer of biotin-fluorescein-dextran labeled BMMs to *Apoe*^−/−^ mice, tissues were stained with 1 mg/mL of streptavidin-A546 for 30 min and nuclei stained using 1 μg/mL of Hoechst for 30 min.

### Statistical Analysis

Statistical analysis was performed using Excel. For comparisons of two groups, student’s t test was used. For comparisons of more than two groups, one-way ANOVA followed by Bonferroni correction was used.

## RESULTS

### LS formation is inhibited in macrophages from Tlr4^−/−^ mice in a MyD88/CD14 dependent manner

To investigate the role of TLR4 in macrophage uptake of agLDL, we examined the interaction of BMMs from *Tlr4*^−/−^ mice and agLDL. We have shown previously that compartments used for degradation of agLDL are formed using F-actin driven membrane protrusions surrounding the aggregate ^11, 14, 20^. We examined F-actin near sites of contact with agLDL in both WT and *Tlr4*^−/−^ BMMs. After a 1 hr incubation with agLDL, an enrichment of F-actin was seen near sites of WT BMM in contact with agLDL (arrow, Figure 1A), but local actin polymerization was greatly reduced in *Tlr4*^−/−^ BMMs (arrow, Figure 1B). To confirm the role of TLR4, we treated cells with monophosphoryl lipid A (MPLA), a TLR4 agonist. WT, but not *Tlr4*^−/−^ BMMs showed an increase in local F-actin staining associated with agLDL upon incubation with MPLA (Figure 1C-D). Further, we treated RAW264.7 macrophages with agLDL for 1 hr and observed TLR4 clustering (a surrogate for TLR4 activation) at the contact site between the macrophage and agLDL (Figure I in the online-only Data Supplement). These data further support a specific role for TLR4 in agLDL induced responses.

**Figure 1.**
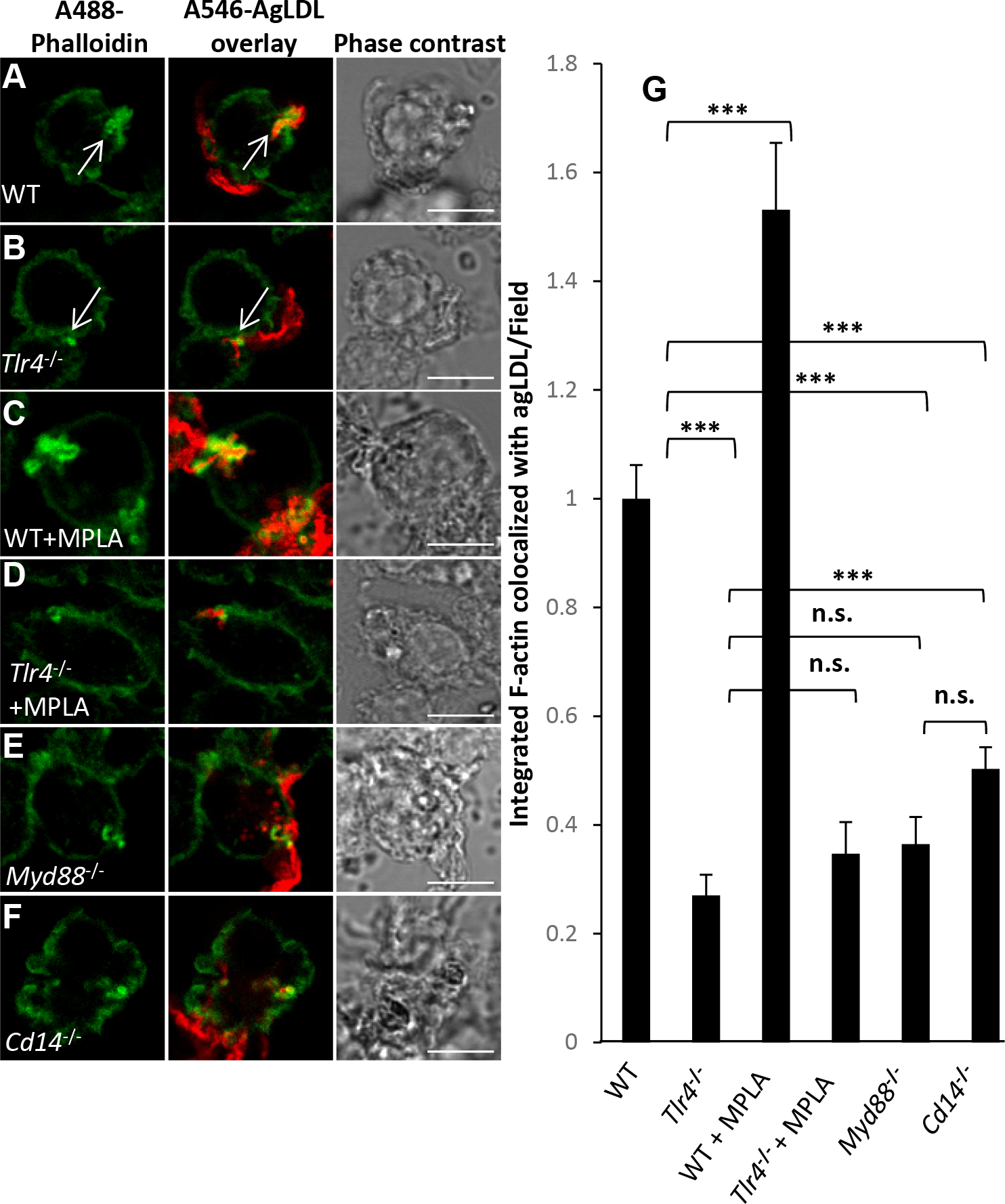
TLR4 regulates actin polymerization at the LS in a MyD88/CD14-dependent manner. Actin polymerization at the LS was analyzed by confocal microscopy of untreated (A) WT and (B) *Tlr4*^−/−^ BMMs, or (C-D) pre-treated with TLR4 agonist MPLA (1 µg/mL) for 1 hr, prior to incubation with A546-agLDL for 1 hr in presence of agonist. (E) *Myd88*^−/−^ and (F) *Cd14*^−/−^ BMMs were also incubated with A546-agLDL for 1 hr. Cells were fixed, and F-actin stained using A488-phalloidin. Arrows highlight areas of macrophage actin polymerization at sites of contact with agLDL. (G) Confocal images were used to quantify F-actin colocalized with agLDL per field. Data were compiled from at least 3 independent experiments per condition. Error bars represent the standard error of the mean (SEM). *** p ≤ 0.001. n.s. not statistically significant (p > 0.05). Scale bars 20 μm.

TLR4 signaling can be broadly divided into MyD88-dependent and MyD88-independent pathways ^25^. Thus, we examined the formation of local F-actin structures at sites of contact with agLDL in *Myd88*^−/−^ BMMs. Compartment formation at sites of contact with agLDL was inhibited in *Myd88*^−/−^ BMMs compared to WT BMMs (Figure 1E). CD14 acts as a co-receptor for TLR4 in the detection of lipopolysaccharide (LPS) and regulates its endocytosis ^26^. Local actin polymerization surrounding agLDL was also inhibited in *Cd14*^−/−^ BMMs compared to WT BMMs (Figure 1F). Quantification of the average local F-actin intensity per field in regions of contact with agLDL revealed a 50-70% decrease in *Tlr4*^−/−^, *Myd88*^−/−^ and *Cd14*^−/−^ BMMs and a 50% increase in MPLA treated WT BMMs (Figure 1G). These results indicate that the formation of extracellular compartments for the catabolism of agLDL occurs, in part, through TLR4 in a MyD88/CD14-dependent fashion.

### Lysosome exocytosis to the LS is inhibited in macrophages from Tlr4^−/−^ mice in a MyD88/CD14-dependent manner

We have previously reported that LS compartments used for extracellular catabolism function as an extracellular hydrolytic organelle due to targeted exocytosis of lysosomes and compartment acidification ^10^. We tested whether lysosomal contents, which include lysosomal acid lipase, a lysosomal hydrolase essential for the degradation of cholesteryl esters, were delivered to points of contact with the aggregate in WT and *Tlr4*^−/−^ BMMs (Figure 2A and B). Biotin-fluorescein-dextran was incubated with BMMs overnight, leading to endocytosis of the dextran and delivery to lysosomes ^10^. Cells were then exposed to streptavidin-A546-labeled agLDL for 90 min. Next, excess biotin was applied to block any remaining free streptavidin on the aggregates. Cells were then fixed and permeabilized to wash away any unbound lysosomal biotin-fluorescein-dextran.

**Figure 2.**
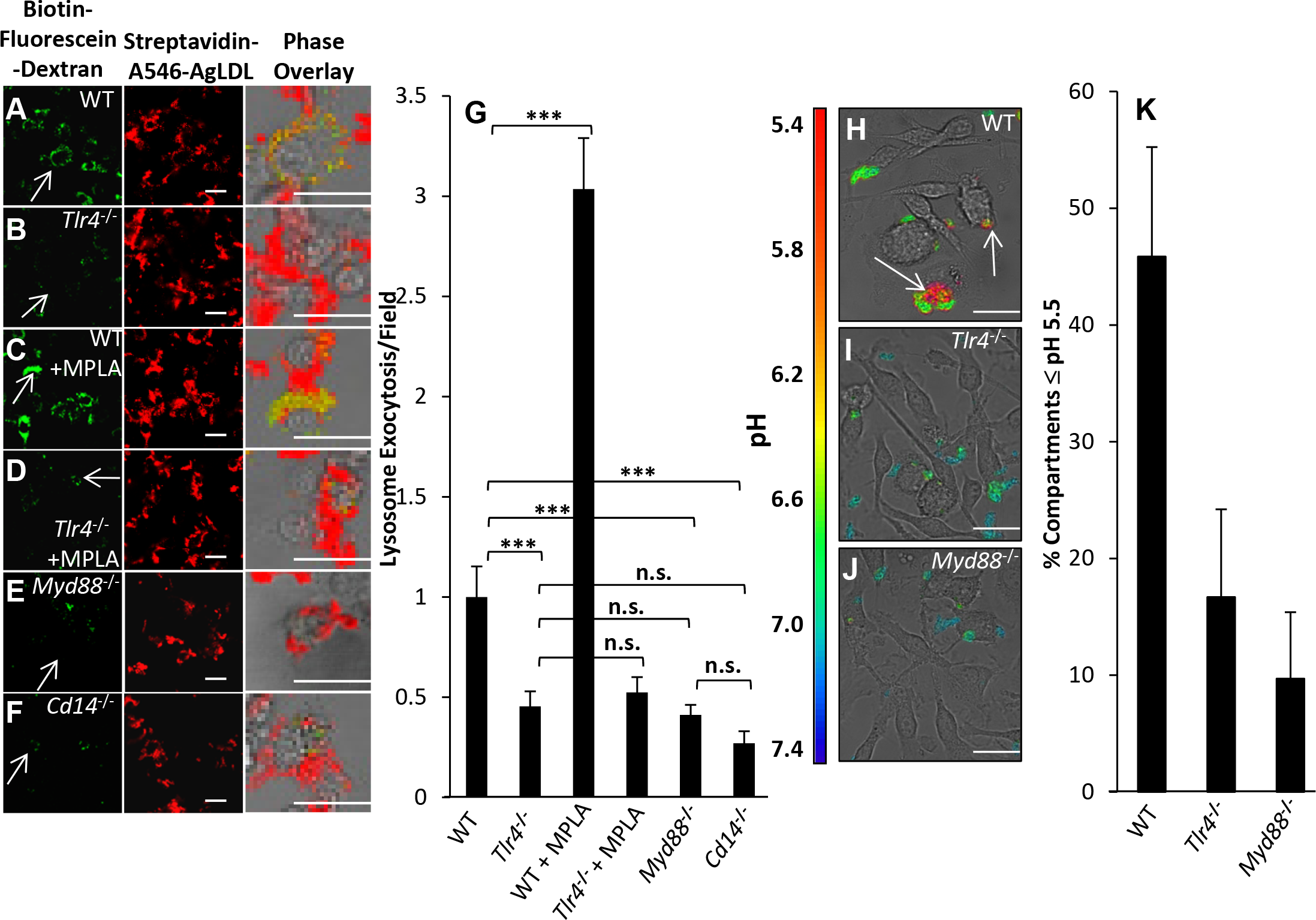
TLR4 regulates lysosome exocytosis at the LS in a MyD88/CD14-dependent manner. (A-G) To assess lysosome exocytosis at the LS, all BMMs were incubated with biotin-fluorescein-dextran overnight and chased in media to dextran load their lysosomes. Subsequently, (A) WT and (B) *Tlr4*^−/−^ BMMs that were left untreated, or (C-D) pre-treated with TLR4 agonist MPLA (1 µg/mL) for 1 hr, were incubated with Streptavidin-A546-agLDL for 90 min in the presence of agonist. (E) *Myd88*^−/−^ and (F) *Cd14*^−/−^ BMMs were also incubated with Streptavidin-A546-agLDL for 90 min. Cells were incubated with an excess of biotin to block unbound streptavidin on agLDL, fixed, permeabilized with triton X-100 and washed extensively to remove biotin-fluorescein-dextran that had not been exocytosed. Cells were then analyzed by confocal microscopy. Arrows highlight areas of macrophage lysosome exocytosis, where lysosomal biotin-fluorescein-dextran has been secreted to the LS and has bound to Streptavidin-A546-agLDL. (G) Confocal images were used to quantify exocytosed biotin-fluorescein-dextran colocalized with Streptavidin-A546-agLDL (lysosome exocytosis) per field. (H-K) Acidification of the LS was assessed by confocal microscopy and ratiometric imaging of (H) WT, (I) *Tlr4*^−/−^ and (J) *Myd88*^−/−^ BMMs treated with CypHer 5E (pH-sensitive) and A488 (pH-insensitive) dual-labeled agLDL for 30 min. (K) Ratiometric images (CypHer 5E/A488) were used to quantify acidification of the LS. Data were compiled from at least 3 independent experiments per condition. Error bars represent the standard error of the mean (SEM). *** p ≤ 0.001. n.s. not statistically significant (p > 0.05). Scale bars 20 μm.

When WT BMMs were in contact with agLDL for 90 min, there was significant deposition of biotin-fluorescein-dextran, due to lysosome exocytosis, onto agLDL (Figure 2A), which was inhibited in *Tlr4*^−/−^ BMMs (Figure 2B). Arrows indicate macrophages, which are shown at higher magnification in the overlay. To confirm the role of TLR4, we treated cells with MPLA. WT BMMs incubated with agLDL in the presence of MPLA showed an increase in lysosome exocytosis (Figure 2C), while *Tlr4*^−/−^ BMMs were unaffected by MPLA treatment (Figure 2D). As shown by quantitative analysis in Figure 2G, TLR4 plays an important role in lysosome exocytosis to the LS. We also found an inhibition in lysosome exocytosis to points of contact with agLDL in *Myd88*^−/−^ and *Cd14*^−/−^ BMMs (Figure 2E-F). Quantification of the amount of lysosome exocytosis shows a 50-70% decrease in *Tlr4*^−/−^, *Myd88*^−/−^ and *Cd14*^−/−^ BMMs compared to WT (Figure 2G). *Tlr4*^−/−^, *Myd88*^−/−^ and *Cd14*^−/−^ BMMs displayed similar lysosome loading of biotin-fluorescein-dextran overnight compared to WT BMMs (Figure II in the online-only Data Supplement).

### Compartment acidification is inhibited in macrophages from Tlr4^−/−^ mice in a MyD88 dependent manner

We have shown previously that the LS becomes acidified through the action of V-ATPase, and this allows lysosomal enzymes delivered to the LS to be active^10^. We examined the role of TLR4 signaling in acidification of extracellular compartments at points of contact with agLDL. To test whether portions of the compartment are acidified, we labeled LDL with CypHer 5E, a pH sensitive fluorophore, and Alexa488, a pH insensitive fluorophore, and performed ratiometric imaging as described previously ^10^. When WT BMMs interacted with the dual labeled agLDL, regions of low pH could be seen at the contact sites (arrows, Figure 2H). Compartment acidification was inhibited in *Tlr4*^−/−^ and *Myd88*^−/−^ BMMs (Figure 2I-J). We quantified the percentage of cells able to acidify to pH 5.5 or lower, as this is the optimal pH for lysosomal acid lipase activity. A reduction in compartment acidification was found in *Tlr4*^−/−^ and *Myd88*^−/−^ BMMs compared to WT BMMs (Figure 2K).

### Foam cell formation in response to agLDL but not monomeric acLDL is inhibited in macrophages from Tlr4^−/−^ mice in a MyD88 dependent manner

Next, we investigated whether the observed inhibition of compartment formation and function affects foam cell formation. After treatment with agLDL for 12 hours, we observed less LipidTOX staining of lipid droplets in *Tlr4*^−/−^ and *Myd88*^−/−^ BMMs than in WT BMMs, indicating decreased levels of intracellular neutral lipid and an inhibition of foam cell formation (Figure 3A-D). This lipid was contained in cytoplasmic lipid droplets, as most of the LipidTOX signal did not colocalize with undegraded LDL in endosomes, and lipid droplets can be visualized in phase contrast images (Figure 3A-C). Further, this inhibition of foam cell formation was specific for agLDL, as WT, *Tlr4*^−/−^ and *Myd88*^−/−^ BMMs treated with acLDL for 12 hours displayed no significant differences in foam cell formation (Figure 3E-H). Taken together, these data show that agLDL causes activation of TLR4 and its immediate downstream effector MyD88 in a CD14 dependent manner. However, TLR4 absence leads to partial reduction of compartment formation, indicating that in addition to TLR4, agLDL likely engages other signaling pathways.

**Figure 3.**
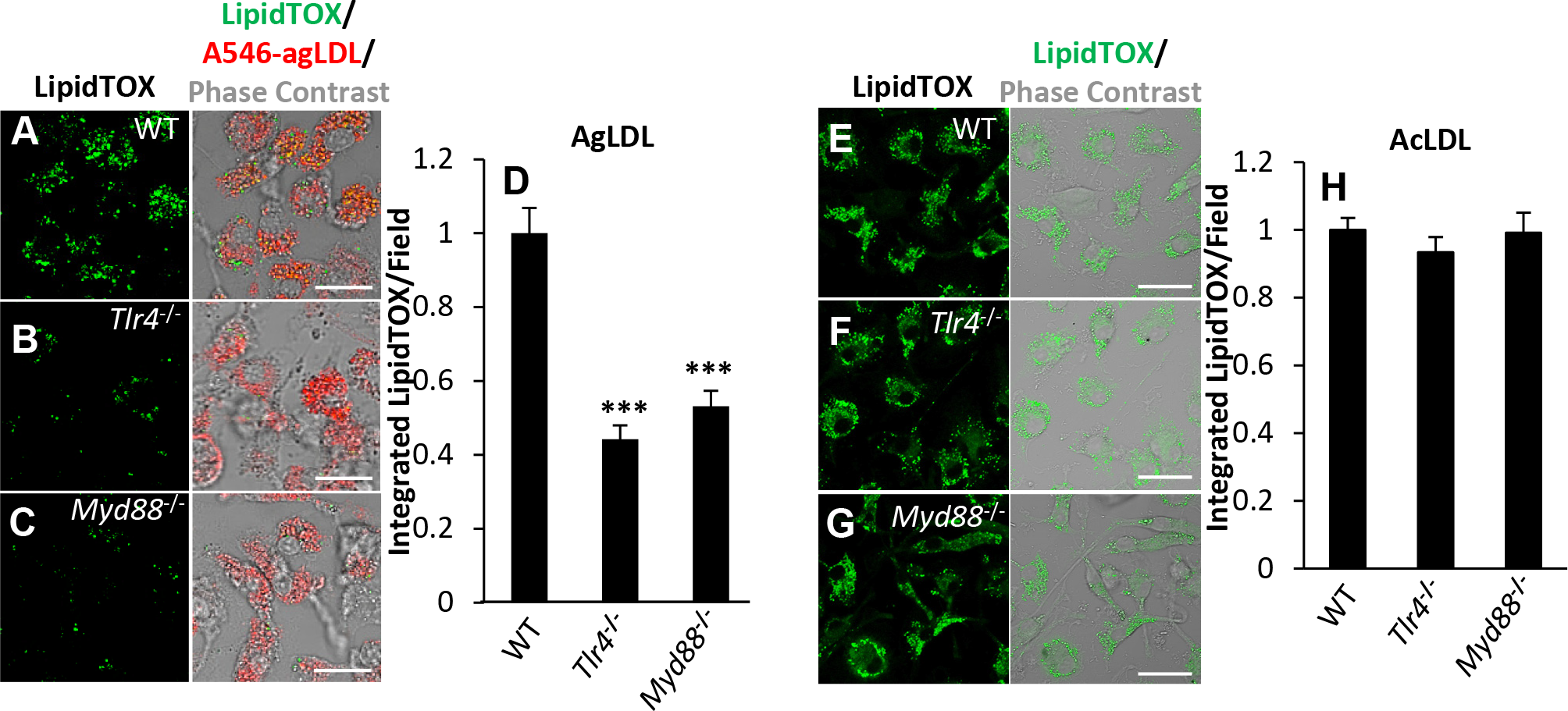
TLR4 and MyD88 regulate foam cell formation in response to agLDL but not acLDL. (A-D) Foam cell formation in response to agLDL was analyzed by confocal microscopy of (A) WT, (B) *Tlr4*^−/−^ and (C) *Myd88*^−/−^ BMMs treated with A546-agLDL for 12 hr, fixed and neutral lipids stained using LipidTOX green. (D) Confocal images were used to quantify LipidTOX intensity per field. (E-H) Foam cell formation in response to acLDL was analyzed by confocal microscopy of (E) WT, (F) *Tlr4*^−/−^ and (G) *Myd88*^−/−^ BMMs treated with acLDL (100 µg/mL) for 12 hr, fixed and neutral lipids stained using LipidTOX green. (H) Confocal images were used to quantify LipidTOX intensity per field. Data were compiled from 3 independent experiments per condition. Error bars represent the standard error of the mean (SEM). *** p ≤ 0.001. n.s. not statistically significant (p > 0.05). Scale bars 20 μm.

### LRP-1 plays a role in lysosome exocytosis to agLDL containing compartments but not local actin polymerization

Absence of TLR4 does not completely abolish agLDL catabolism (Figure 3), indicating the involvement of additional receptors. Several reports have demonstrated the involvement of LRP-1 in macrophage uptake of agLDL ^27, 28^. LRP-1 is the major LRP expressed in macrophages and has been implicated to play a role in atherosclerosis ^29^. To see if LRP-1 plays a role in compartment formation and function, we inhibited LRP-1 with receptor-associated protein (RAP) and assessed actin polymerization and lysosome exocytosis as in Figures 1 and 2. We had previously shown that inhibition of LRP-1 does not impair actin polymerization at points of contact with the aggregate in WT BMM ^14^, and here we found the same to be true for *Tlr4*^−/−^ BMM (Figure III A in the online-only Data Supplement). When we examined lysosome exocytosis to the compartment, we found decreased exocytosis in WT BMMs treated with RAP compared to untreated controls (Figure III B in the online-only Data Supplement). We also examined inhibition of LRP-1 in *Tlr4*^−/−^ BMMs. We found a modest decrease in lysosome exocytosis of *Tlr4*^−/−^ BMMs treated with RAP (Figure III B in the online-only Data Supplement), indicating that TLR4 is not necessary for the LRP-1 agLDL interaction.

Next we examined the involvement of several receptors, LDL receptor (LDLr), SCARA1 and CD36, which are known to play a role in the uptake of various species of mLDL. CD36, a scavenger receptor important for the uptake of oxidized LDL also participates in TLR4 signaling triggered by LPS ^30^. When *Ldlr*^−/−^, *Scara1*^−/−^, or *Cd36*^−/−^ BMMs were incubated with agLDL, no difference in compartment formation (local actin polymerization) (Figure III C in the online-only Data Supplement), lysosome exocytosis (Figure III D in the online-only Data Supplement) or foam cell formation (Figure III E in the online-only Data Supplement) was seen. These results demonstrate the lack of involvement of these scavenger receptors in agLDL catabolism by macrophages, clearly distinguishing this mechanism of uptake and foam cell formation from those involving receptor mediated mLDL or oxidized LDL uptake.

### Macrophage catabolism of agLDL requires spleen tyrosine kinase

SYK constitutively binds to the cytoplasmic toll/interleukin-1 receptor domain of TLR4 ^19^ and plays a prominent role in TLR4 signaling in response to minimally modified LDL ^19, 31^. Depletion of SYK in J774 cells was achieved using siRNA treatment, and SYK protein levels were reduced by 80% compared to control siRNA treated cells (Figure 4A-B). We observed a significant decrease in actin polymerization (Figure 4C, Figure IV A-B in the online-only Data Supplement) and lysosome exocytosis (Figure 4D, Figure IV C-H in the online-only Data Supplement) at sites of contact with agLDL. SYK knockdown did not impair lysosome loading of biotin-fluorescein-dextran (Figure IV I in the online-only Data Supplement). These results were confirmed using a specific inhibitor of SYK, BAY 61-3606. When WT BMMs were treated with BAY 61-3606, a large reduction in local actin polymerization was seen at sites of contact with agLDL (Figure 4E, Figure IV J-K in the online-only Data Supplement). Surprisingly, when *Tlr4*^−/−^ and *Myd88*^−/−^ BMMs were treated with BAY 61-3606, F-actin mediated compartment formation was further reduced (Figure 4E, Figure IV L-O in the online-only Data Supplement). This indicates that in addition to TLR4, SYK is interacting with another currently unidentified activation pathway. Likewise, lysosome exocytosis was strongly inhibited upon cellular treatment with BAY 61-3606 in WT, *Tlr4*^−/−^ and *Myd88*^−/−^ BMMs (Figure 4F, Figure IV P-U in the online-only Data Supplement).

**Figure 4.**
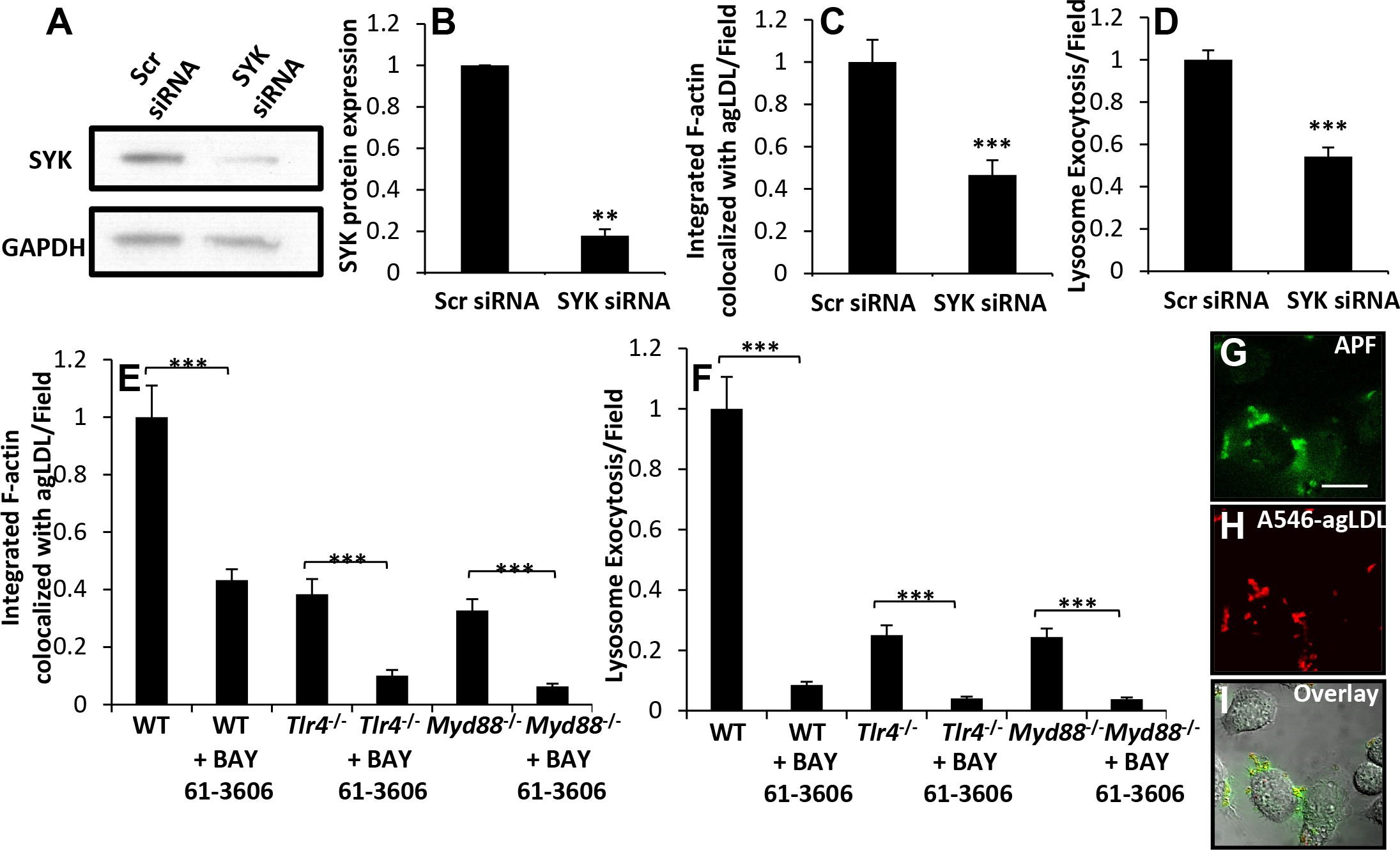
SYK promotes actin polymerization and lysosome exocytosis at the LS. (A-D) J774 cells were treated with scrambled (Scr) control or SYK siRNA for 48 hr. SYK protein levels in Scr and SYK siRNA treated cells were (A) analyzed by immunoblotting and (B) quantified from several independent experiments. GAPDH was used as a loading control. (C) Actin polymerization at the LS was analyzed by confocal microscopy of Scr and SYK siRNA treated J774 cells incubated with A546-agLDL for 1 hr. Cells were fixed, and F-actin stained using A488-phalloidin, prior to analysis. Confocal images were used to quantify F-actin colocalized with agLDL per field. (D) Lysosome exocytosis at the LS was analyzed by confocal microscopy. Scr and SYK siRNA treated J774 cells were incubated with biotin-fluorescein-dextran overnight and chased in media to dextran load their lysosomes. Subsequently, cells were incubated with Streptavidin-A546-agLDL for 90 min. Cells were incubated with an excess of biotin, fixed, permeabilized with triton X-100, washed extensively and analyzed. Confocal images were used to quantify exocytosed biotin-fluorescein-dextran colocalized with Streptavidin-A546-agLDL (lysosome exocytosis) per field. (E) Actin polymerization at the LS was analyzed by confocal microscopy of WT, *Tlr4*^−/−^ and *Myd88*^−/−^ BMMs with treatment using SYK inhibitor BAY 61-3606 (5 μM) or DMSO control for 1 hr prior to treatment with A546-agLDL for 1 hr in the presence or absence of inhibitor. Cells were fixed, and F-actin stained using A488-phalloidin, prior to analysis. Confocal images were used to quantify F-actin colocalized with agLDL per field. (F) Lysosome exocytosis at the LS was analyzed by confocal microscopy. WT, *Tlr4*^−/−^ and *Myd88*^−/−^ BMMs were incubated overnight with biotin-fluorescein-dextran to load macrophage lysosomes and chased in media. Subsequently, cells were treated SYK inhibitor BAY 61-3606 (5 μM) or DMSO control for 1 hr prior to treatment with Streptavidin-A546-agLDL for 90 min in the presence or absence of inhibitor. Cells were incubated with an excess of biotin, fixed, permeabilized with triton X-100, washed extensively and analyzed. Confocal images were used to quantify biotin-fluorescein-dextran intensity colocalized with Streptavidin-A546-agLDL (lysosome exocytosis) per field. Data were compiled from at least 3 independent experiments per condition, and error bars represent the SEM. ** p ≤ 0.01, *** p ≤ 0.001. (G-I) J774 cells were treated with A546-agLDL for 30 min prior to addition of 15 μM APF (ROS indicator) and live cell imaging was performed. Scale bar 20 μm.

The SYK/TLR4 complex has been shown to play a role in macrophage generation of reactive oxygen species (ROS) in response to minimally modified LDL ^31^. Thus, we looked for evidence of ROS generation in the extracellular compartments used for catabolism of agLDL. ROS generation was observed (green, Figure 4G-I) colocalized with agLDL as well as diffusing away from the LS, using the ROS indicator aminophenyl fluorescein (APF).

### Inhibition of PI3K greatly reduces macrophage catabolism of agLDL

Activated SYK can directly bind to PI3K ^32^. To assess the role of PI3K in macrophage catabolism of agLDL, we treated J774 cells with LY294002, an inhibitor of both class I and class II PI3K ^33^. Inhibition of PI3K abolishes F-actin assembly at sites of contact with agLDL (Figure 5A). Class IA PI3K are known to bind to phosphorylated tyrosine ^34^, and can have important roles in different cell types. To further characterize the role of PI3K in agLDL degradation in macrophages, we investigated the role of class I catalytic subunits in macrophage catabolism of agLDL. Treatment of J774 cells with various inhibitors specific for the catalytic subunits of Class I PI3K ^33, 35^ shows that inhibitors of four subunits (p110α, p110β, p110δ and p110γ) reduce the formation of the local F-actin rich structures used to form compartments for agLDL degradation (Figure 5A, Figure V A-F in the online-only Data Supplement). We also examined the role of PI3K in lysosome exocytosis to compartments used for degradation of agLDL. LY294002 treatment completely abolished lysosome exocytosis, indicating that PI3K is a key signaling molecule in macrophage degradation of agLDL (Figure 5B, Figure V G-L in the online-only Data Supplement). In contrast to effects on actin polymerization, only two of the four Class I PI3K subunits were found to be involved in lysosome exocytosis to agLDL containing compartments. Inhibition of the Class IA p110β and p110δ subunits resulted in a decrease in lysosome exocytosis (Figure 5B). The Class IA p110α and the Class IB p110γ subunits do not play a role in lysosome exocytosis during macrophage catabolism of agLDL (Figure 5B, Figure V G-L in the online-only Data Supplement). Inhibition of the p110β subunit resulted in the largest reduction in macrophage catabolism of agLDL. This subunit also plays a key role in osteoclast bone remodeling, a related process in which degradation occurs extracellularly ^36^.

**Figure 5.**
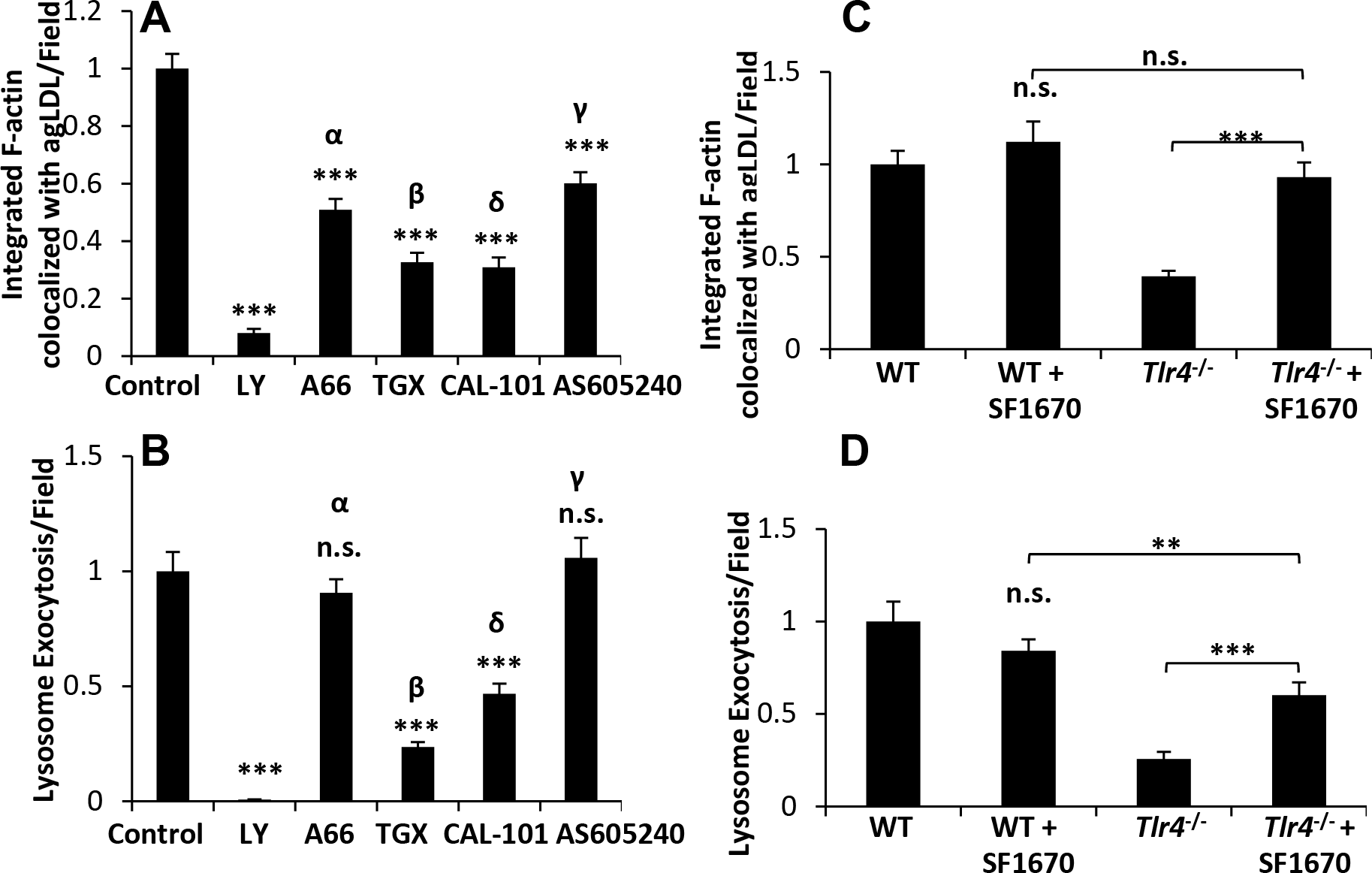
PI3K regulates actin polymerization and lysosome exocytosis at the LS in a TLR4-dependent manner. (A) Actin polymerization to the LS was analyzed by confocal microscopy of J774 cells pre-treated with DMSO (control), LY294002 (50 μM), p110α inhibitor A66 (8 μM), p110β inhibitor TGX-221 (2 μM), p110δ inhibitor CAL-101 (2 μM) and p110γ inhibitor AS-605240 (2 μM) for 1 hr prior to treatment with A546-agLDL for 1 hr in the presence or absence of inhibitors. Cells were fixed, and F-actin was stained using A488-phalloidin, prior to analysis. Confocal images were used to quantify F-actin colocalized with agLDL per field. (B) Lysosome exocytosis at the LS was analyzed by confocal microscopy. J774 cells were incubated with biotin-fluorescein-dextran overnight and chased in media to dextran load their lysosomes. Cells were pre-treated with the same inhibitors as in (A) prior to treatment with Streptavidin-A546-agLDL for 90 min in the presence or absence of inhibitors. Cells were incubated with an excess of biotin, fixed, permeabilized with triton X-100, washed extensively and analyzed. Confocal images were used to quantify exocytosed biotin-fluorescein-dextran colocalized with Streptavidin-A546-agLDL (lysosome exocytosis) per field. (E) Actin polymerization to the LS in WT and *Tlr4*^−/−^ BMMs pre-treated with DMSO control or PTEN inhibitor SF1670 (1 μM) for 1 hr prior to treatment with A546-agLDL for 1 hr in the presence or absence of inhibitors was assessed by confocal microscopy and quantified per field. (F) Lysosome exocytosis at the LS in WT and *Tlr4*^−/−^ BMMs pre-treated under the same conditions as (E) prior to treatment with Streptavidin-A546-agLDL for 90 min in the presence of inhibitors was assessed by confocal microscopy and quantified per field. Data were compiled from at least 3 independent experiments per condition. Error bars represent the SEM. ** p ≤ 0.01, *** p ≤ 0.001. n.s. not statistically significant.

### TLR4 promotes macrophage catabolism of agLDL in a PI3K dependent manner

To investigate the link between TLR4/MyD88 and PI3K, we inhibited the PI(3,4,5)P_3_ phosphatase PTEN (phosphatase and tensin homologue deleted on chromosome 10). When *Tlr4*^−/−^ BMMs were treated with 1 μM SF1670, a PTEN inhibitor used previously ^37^, both local actin polymerization (Figure 5C, Figure V M-P in the online-only Data Supplement) and lysosome exocytosis (Figure 5D, Figure V Q-T in the online-only Data Supplement) were increased compared to untreated cells. No difference was observed between WT BMMs treated with SF1670 and control cells (Figure 5C-D). These results indicate that TLR4 is upstream of PI3K and that the effects of TLR4 removal can be reversed by inhibition of PTEN.

### PI3K signals through Akt during macrophage catabolism of agLDL

The activation of the serine/threonine kinase Akt is mainly mediated by PI3K generation of PIP_3_, which recruits Akt and other PH domain-containing proteins to the plasma membrane ^34^. To test the involvement of Akt in the degradation of agLDL, J774 cells were treated with agLDL in the presence or absence of 4 μM Akti1/2, which inhibits both Akt1 and Akt2. Both compartment formation (local actin polymerization) (Figure 6A, Figure VI A-B in the online-only Data Supplement) and lysosome exocytosis to the compartment (Figure 6B, Figure VI C-D in the online-only Data Supplement) were reduced in cells treated with Akti1/2. The mammalian serine/theronine Akt kinases comprise three closely related isoforms: Akt1, Akt2 and Akt3. Akt1 and Akt2 are both highly expressed in macrophages ^38^. When *Akt1*^−/−^ and *Akt2*^−/−^ BMMs were incubated with A546-agLDL for 1 hour, actin polymerization at sites of contact with the aggregate was reduced in *Akt2*^−/−^, but not *Akt1*^−/−^ macrophages (Figure 6C, Figure VI E-G in the online-only Data Supplement). However, absence of either isoform led to reduced lysosome exocytosis to agLDL containing compartments (Figure 6D, Figure VI H-J in the online-only Data Supplement), but did not affect lysosome loading of biotin-fluorescein-dextran (Figure VI K in the online-only Data Supplement). These results suggest that Akt1 plays a role in lysosome exocytosis but not compartment formation, while Akt2 is required for both processes during agLDL catabolism.

**Figure 6.**
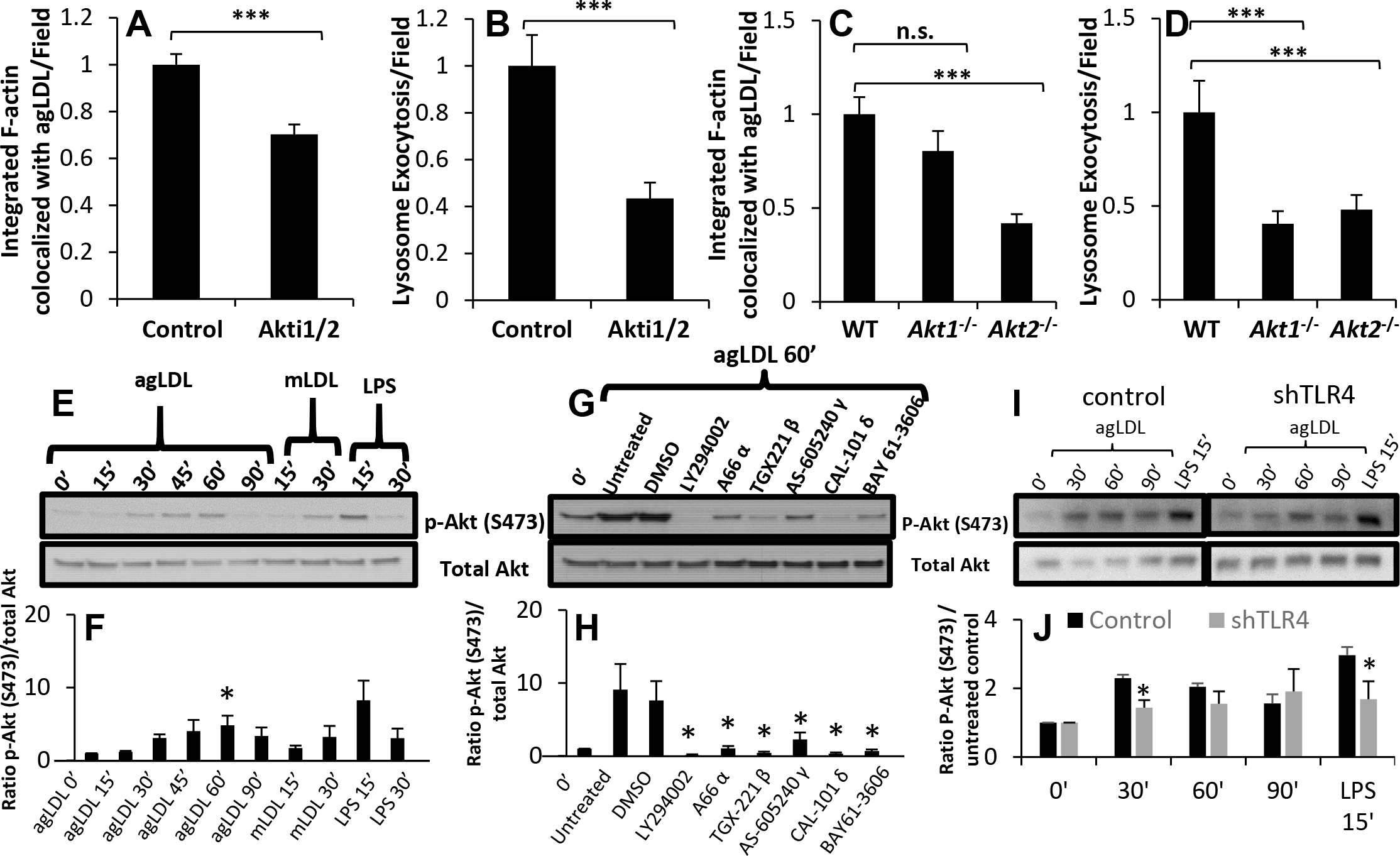
AgLDL-stimulated Akt activation regulates actin polymerization and lysosome exocytosis at the LS and occurs in a PI3K/SYK dependent manner. (A) Actin polymerization at the LS was analyzed by confocal microscopy of J774 cells pre-treated with DMSO (control) or Akt inhibitor Akti1/2 (4 μM) for 1 hr prior to treatment with A546-agLDL for 1 hr in the presence or absence of inhibitor. Cells were fixed, and F-actin stained using A488-phalloidin, prior to analysis. Confocal images were used to quantify F-actin colocalized with agLDL per field. (B) Lysosome exocytosis at the LS was analyzed by confocal microscopy. J774 cells were incubated with biotin-fluorescein-dextran overnight and chased in media to dextran load their lysosomes. Cells were pre-treated with the same inhibitors as in (A) prior to treatment with Streptavidin-A546-agLDL for 90 min in the presence or absence of inhibitors. Cells were incubated with an excess of biotin, fixed, permeabilized with triton X-100, washed extensively and analyzed. Confocal images were used to quantify exocytosed biotin-fluorescein-dextran colocalized with Streptavidin-A546-agLDL (lysosome exocytosis) per field. (C) Actin polymerization at the LS was analyzed by confocal microscopy of WT, *Akt1*^−/−^ and *Akt2*^−/−^ BMM treated with A546-agLDL for 1 hr and quantified per field. (D) Lysosome exocytosis at the LS was assessed by confocal microscopy of WT, *Akt1*^−/−^ and *Akt2*^−/−^ BMM treated with Streptavidin-A546-agLDL for 90 min and quantified per field. (E) J774 cells were treated with agLDL (1 mg/mL), mLDL (1 mg/mL) or LPS (100 ng/mL) for indicated periods of time, prior to lysis and immunoblot analysis. (F) Ratios of phosphorylated (S473) to total Akt protein were obtained by densitometry analysis. (G) J774 cells were pre-treated for 1 hr with various inhibitors (LY294002 50 μM, A66 8 μM, TGX-221 2 μM, AS-605240 2 μM, CAL-101 2 μM, and BAY 61-3606 5 μM), left untreated or treated with DMSO control, and subsequently incubated with agLDL (1 mg/mL) for 1 hr in the presence or absence of inhibitors prior to lysis and immunoblot analysis. (H) Ratios of phosphorylated (S473) to total Akt protein were obtained by densitometry analysis. (I) J774 cells stably transfected with control shRNA or TLR4 specific shRNA (shTLR4) were treated with agLDL (1 mg/mL) for indicated periods of time prior to lysis and immunoblot analysis. (J) Ratios of phosphorylated (S473) to total Akt protein were obtained by densitometry analysis.. Data were compiled from 3 independent experiments. Error bars represent the SEM. * p < 0.05, ** < 0.01, *** p ≤ 0.001. n.s. not statistically significant.

Having seen an effect by inhibition or knockout of Akt1/2, we examined whether agLDL could induce Akt activation. J774 cells were incubated with agLDL for 15 - 90 min, and Akt phosphorylation at Ser473 and Thr308 was analyzed. LPS served as a positive control. Treatment of J774 cells with mLDL for 15 and 30 min was also performed to compare agLDL and mLDL responses. Incubation of cells with agLDL induced phosphorylation at the Ser473 site (Figure 6E), demonstrating that agLDL induces Akt activation. Quantification of the phosphorylation indicates that agLDL treatment induces peak activation at 45-60 min for the Ser473 site (Figure 6F) and 45 min for the Thr308 site (Figure VII B in the online-only Data Supplement). Incubation with mLDL also modestly increased phosphorylation of Akt at the Ser473 site (Figure 6E). Similar results were observed for the Thr308 site (Figure VII A in the online-only Data Supplement). Modest Akt activation has been reported in monocytes and BMMs incubated with mLDL ^39, 40^. LPS increased Akt phosphorylation at both sites effectively (Figure 6F and Figure VII B in the online-only Data Supplement). These data suggest that agLDL can activate Akt.

Inhibition of SYK, PI3K and the PI3K catalytic subunits p110α, p110β, p110γ and p110δ reduce actin polymerization and lysosome exocytosis to the compartment (Figures 4 and 5). We tested whether these effects could be explained by diminished Akt activation. J774 cells were treated with inhibitors for these proteins and incubated with agLDL for 1 hr. Quantification of the magnitude of phosphorylation of Akt at Ser 473 relative to total Akt revealed that treatment with LY294002, a PI3K inhibitor completed abolished Akt phosphorylation (Figure 6G-H). Consistent with their effects on local actin polymerization and lysosome exocytosis, treatment with p110α or p110γ specific inhibitors diminished Akt phosphorylation to a limited extent, while p110β and p110δ inhibitors were more effective at reducing Akt phosphorylation stimulated by agLDL (Figure 6G-H). Inhibition of SYK by BAY 61-3606 inhibited Akt phosphorylation induced by agLDL treatment at the Ser473 site (Figure 6G-H). Again, similar results were obtained for the Thr308 site (Figure VII C-D in the online-only Data Supplement). These data indicate that SYK is able to act upstream of Akt in agLDL induced signaling and that PI3K plays an essential role in Akt phosphorylation and activation in response to agLDL treatment.

As reported previously, downregulation of TLR4 in J774 cells using shRNA reduced surface levels of TLR4 by 60% compared to control shRNA cells ^19^. Using these cells, downregulation of TLR4 delays activation of Akt compared to control cells (Figure 6I-J). When TLR4 was reduced, Akt phosphorylation at the Ser473 site was significantly reduced after 30 min agLDL treatment (Figure 6I-J), though this was a partial effect. This partial reduction in Akt activation may be explained by residual TLR4 expression or may be due to other activation pathways. Consistent with our previous data, these data indicate that TLR4 plays a role in downstream activation of Akt in response to agLDL.

### *Tlr4*^−/−^ macrophages isolated from murine atherosclerotic plaques exhibit inhibition of lipid accumulation compared with WT macrophages

TLR4 signaling is known to be involved in atherosclerosis, but few *in vivo* studies have focused on the role of TLR4 in macrophage foam cell formation. One study found that hyperlipidemic TLR4 whole body knockout mice accumulate less lipid in the aortic arch, though alterations in smooth muscle cells were implicated to be the cause of this ^41^. TLR4 whole body knockout is likely to alter functions in many different cells types. To specifically examine the role of TLR4 in macrophage lipid accumulation *in vivo*, we used a bone marrow transplantation strategy. Following irradiation, *Ldlr*^−/−^ mice were reconstituted with marrows from either WT or *Tlr4*^−/−^ mice and placed on a high fat diet (HFD) for 16 weeks. Cell suspensions were prepared from aortas, and cells were cytospun onto glass slides. Cells were stained with an antibody to F4/80 to identify macrophages and LipidTOX green to quantify neutral lipid content (Figure 7A-J).

**Figure 7.**
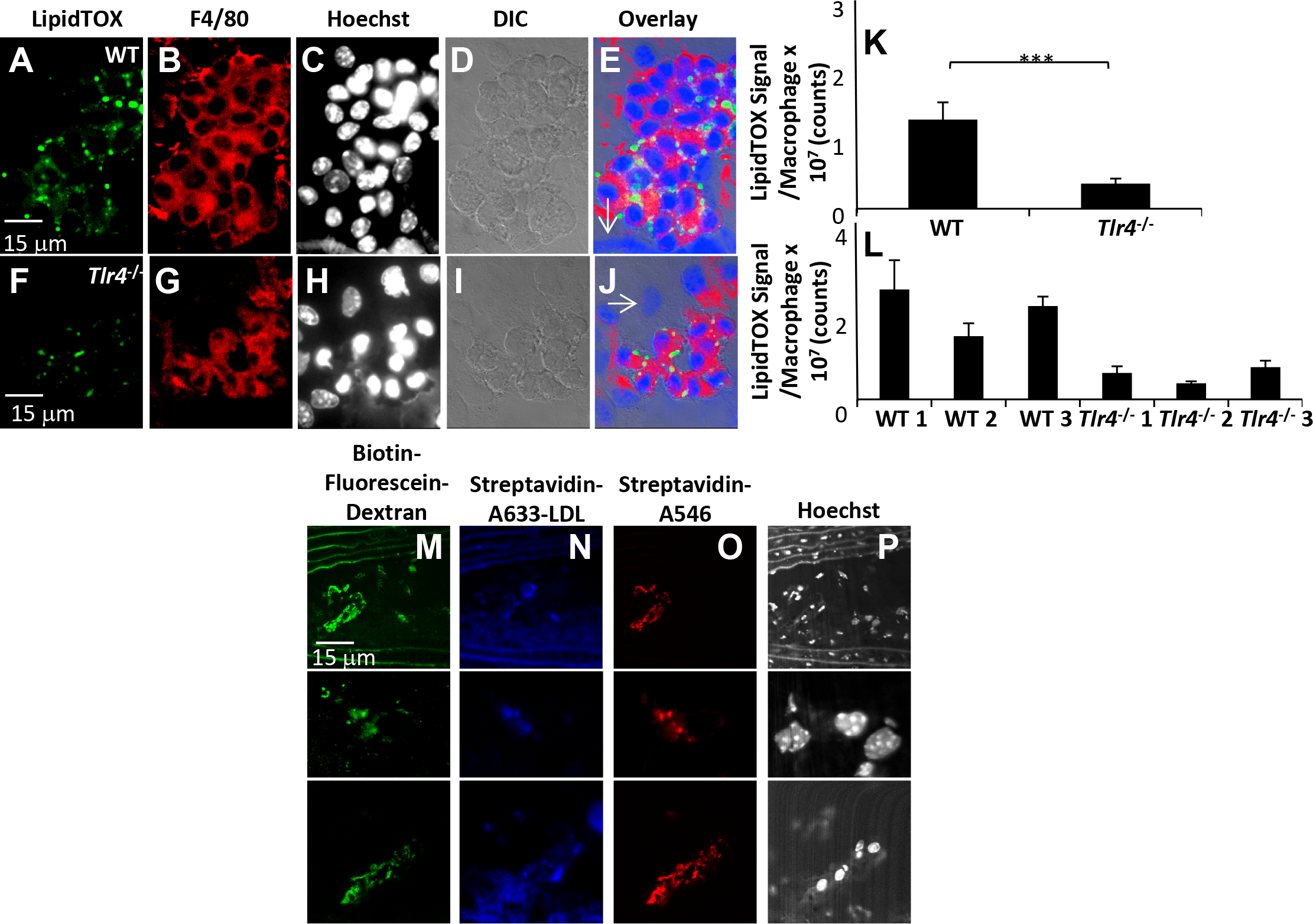
Macrophage deficiency of TLR4 protects against cholesterol uptake and foam cell formation *in vivo* during atherosclerosis. *Ldlr*^−/−^ mice were gamma irradiated and reconstituted with WT (A-E) or *Tlr4*^−/−^ (F-J) marrows. Mice received a HFD for 16 weeks and were sacrificed. Aortas were digested, and single cell suspensions were centrifuged onto coverslips. Cells were stained to detect neutral lipid (LipidTOX green) (A, F), macrophages (F4/80) (B, G) and Hoechst (nuclei) (C, H), analyzed by confocal microscopy and LipidTOX fluorescence per macrophage from WT or *Tlr4*^−/−^ reconstituted mice was quantified. Arrows denote cells negative for F4/80 staining. Data shown are compiled from 6 mice (3 WT and 3 *Tlr4*^−/−^ reconstituted mice) (K) and quantification split per mouse (L). Error bars represent the SEM. *** p ≤0.001. (M-O) *In vivo* example of a macrophage using a lysosomal synapse to degrade agLDL. *Apoe*^−/−^ - mice were placed on a HFD for 24 weeks. Animals were sacrificed, and their aortas removed, fixed, sectioned and stained with filipin (green), to show cholesterol, and an F4/80 antibody (red) to show macrophages. Plaques were examined using confocal microscopy. (M-P) *Apoe*^−/−^ mice on a HFD were injected with streptavidin-A633-LDL 1 day prior to injection with BMMs with their lysosomes loaded with biotin-fluorescein-dextran. 3 days after adoptive transfer, mice were sacrificed, and the aortas were harvested, sectioned, incubated with streptavidin-A546 and examined by confocal microscopy. (M) Several examples of adoptively transferred BMMs that have exocytosed lysosomal contents (biotin-fluorescein-dextran) at points of contact with injected streptavidin-A633-LDL (N). Streptavidin-A546 labeling (O) is used to confirm the specificity of the interaction. Nuclear staining is also shown (P).

LipidTOX signal was quantified for WT and *Tlr4*^−/−^ macrophages isolated from the plaques of *Ldlr*^−/−^ mice (Figure 7K-L). *Tlr4*^−/−^ macrophages contained less neutral lipid than WT macrophages, indicative of an inhibition of foam cell formation. These results demonstrate that our cell culture findings examining degradation of agLDL using the LS mirror *in vivo* patterns of macrophage lipid accumulation.

We have observed compartment formation in atherosclerotic plaques previously ^20^. To confirm that these extracellular compartments are used for catabolism, we looked for evidence of lysosome exocytosis to agLDL containing compartments in a mouse model of atherosclerosis. Streptavidin-A633-LDL was introduced by tail vein injection into *Apoe*^/-^ mice that had been on the HFD for 24 weeks. One day after the LDL injection, cultured BMMs with lysosomes loaded with biotin-fluorescein-dextran were adoptively transferred to the mice. Mice were euthanized 3 days after the adoptive transfer, and aortic sections were stained with streptavidin-A546 (to confirm that the observed areas of green fluorescence were from biotin-fluorescein-dextran) and examined by confocal microscopy to look for *in vivo* evidence of the delivery of lysosomal contents to points of contact with the aggregate.

We found evidence of the delivery of lysosomal contents to points of contact with the aggregate. Figure 7M shows several examples of lysosome exocytosis to a LS acquired from a plaque in the lesser curvature of the aortic arch of an *Apoe*^−/−^ mouse that underwent injection of streptavidin-A633-LDL and adoptive transfer of macrophages with lysosomes loaded with biotin-fluorescein-dextran. Also shown are the corresponding distribution of the injected streptavidin-A633-LDL (Figure 7N), the streptavidin-A546 labeling, which serves to confirm the specificity of the interaction, (Figure 7O) and nuclear staining (Figure 7P). *In vivo*, we have observed hundreds of cells exhibiting exocytosis to structures similar in morphology to those observed in cell culture in many mice (Figure VIII in the online-only Data Supplement). These results suggest that *in vivo*, macrophages use an extracellular compartment containing secreted lysosomal enzymes to degrade agLDL.

## DISCUSSION

We have described a novel mechanism of macrophage lipid accumulation in which agLDL, the predominant form of LDL found in atherosclerotic plaques, is involved in activation of TLR4. In a process that we have called “exophagy”, macrophages form an extracellular compartment supported by F-actin that surrounds agLDL ^11, 14^, and lysosomes are secreted into these compartments, which are acidified by V-ATPase ^10^. Initial catabolism of the aggregate occurs in these extracellular, acidic, lytic compartments, ultimately resulting in foam cell formation ^10^. Herein we show that TLR4 is required to initiate the robust cytoskeletal rearrangements and lysosome exocytosis that lead to catabolism of agLDL. We also demonstrate involvement of LRP-1, MyD88, SYK, PI3K and Akt in this process (illustrated in the graphic abstract). We show that both *in vitro* and *in vivo*, macrophages lacking TLR4 are protected against internalization of free cholesterol and ensuing foam cell formation and provide *in vivo* evidence that macrophages use an extracellular compartment for the degradation of agLDL. Our findings provide a link between hyperlipidemia and a proinflammatory signaling cascade that is associated with the pathogenesis of atherosclerosis.

An interesting question raised by this work is how TLR4 is activated by agLDL. A recent study demonstrated that a component of oxidized LDL, an oxidized cholesteryl arachidonate with bicyclic endoperoxide and hydroperoxide groups can activate a TLR4/SYK signaling pathway and is present in human plasma and in human atherosclerotic lesions ^42^. It seems likely that this molecule would also play a role in agLDL stimulation of TLR4 signaling shown in this study. Other studies have looked at TLR4 ligand LPS, which contains acylated hydroxyl saturated fatty acids. Removal of these results in a loss of LPS-induced inflammation ^43^. This attribute of LPS led to the hypothesis that saturated fatty acids could be natural ligands for TLR4, a theory that has been confirmed by several groups but remains controversial ^44, 45^. However, studies have also shown activation of TLR4 through changes in the lipid content of the plasma membrane ^46^, which would occur during extracellular catabolism of agLDL. Interestingly, inhibition of ABC transporter mediated cholesterol efflux heightens signaling through TLRs ^47, 48^, indicating that elevated plasma membrane cholesterol may enhance uptake of agLDL by further activating TLR4, resulting in a positive feedback loop. Future studies will investigate whether agLDL directly binds to TLR4 or if activation occurs through changes in membrane lipid content or both.

The observation that *Tlr4*^−/−^ BMMs exhibit a reduction in agLDL uptake, while treatment with a SYK or PI3K inhibitor almost completely abolishes aggregate degradation, points to the involvement of additional signaling pathways. Integrins represent a class of surface receptors that are likely candidates. SYK is known to be activated during integrin signaling and also to bind to β-integrins ^49^, which are important in the formation and function of the extracellular lytic compartment that osteoclasts use for degradation of bone ^50^. The involvement of the PI3K Class IB catalytic subunit p110γ in the local F-actin polymerization used for compartment formation may indicate participation of a G-protein coupled receptor (GPCR) as p110γ is known to be activated downstream of GPCRs ^34^.

Numerous animal studies have been conducted to assess the effect of the receptors and signaling cascades examined herein on atherosclerosis ^15, 17, 51–53^. In particular, foam cell formation in early lesions of *Tlr4*^−/−^*Apoe*^−/−^ mice is reduced by 70-80% compared with *Apoe*^−/−^ controls ^41^. However, production of several chemokines is concomitantly reduced in the absence of TLR4, so monocyte/macrophage trafficking to the plaque is diminished ^15^. In these animals, alterations in macrophage plaque recruitment may explain the observed reduction in foam cell formation rather than a direct inhibition of macrophage lipid accumulation ^15^. Thus, while animal studies provide rich information regarding the role of specific moieties in atherosclerosis, it is a multifaceted disease involving many processes and cells types. This is particularly obvious when we compare our data suggesting that macrophage TLR4 activation by agLDL depends on CD14, with data showing that global CD14 deficiency does not improve atherosclerosis in mice ^17^. Many cell types that express CD14 regulate the pathogenesis of atherosclerosis, so the results need not be at odds. Further, Bjorkbacka et al. confined their examination to the aortic sinus, which may also have masked a role for CD14 elsewhere ^54^.

Limitations in pinpointing the mechanistic basis of effects observed in animal studies highlight the need to couple this work with basic cell biology research. To this end, we investigated a macrophage signaling cascade that occurs during agLDL induced foam cell formation. Our data lead to a more precise understanding of the relationship between atherosclerosis and a maladaptive sterile inflammatory response and provide several novel targets that can be manipulated to assess the relevance of this mechanism of macrophage foam cell formation *in vivo*.

## Supporting information

Supplemental Figures I - VIII

## Abbreviations

agLDL: aggregated LDL
BMM: bone marrow-derived macrophage
F-actin: filamentous actin
FBS: fetal bovine serum
LDL: low-density lipoprotein
LS: lysosomal synapse
WT: wild-type

## ACKNOWLEDGEMENTS

We would like to thank Morris Burnbaum (University of Pennsylvania) for providing *Akt1*^−/−^ and *Akt2*^−/−^ femurs. We thank Aiho Ding (Weill Cornell Medical College) for providing *Tlr4*^−/−^ and *Myd88*^−/−^ femurs for preliminary experiments. We thank Yuri Miller and Soo-Ho Choi (University of California, San Diego) for control and TLR4 shRNA expressing J774 cells.

## HIGHLIGHTS

- Macrophages use a TLR4/MyD88/PI3K/SYK/Akt-dependent signaling pathway to form the LS to degrade agLDL and form foam cells
- Macrophage-specific deficiency of TLR4 protects macrophages from lipid accumulation during atherosclerosis
- Lysosome exocytosis by macrophages occurs *in vivo* during atherosclerosis to the LS to degrade agLDL
- The mechanism of agLDL degradation and uptake can be distinguished from monomeric or oxidized LDL uptake

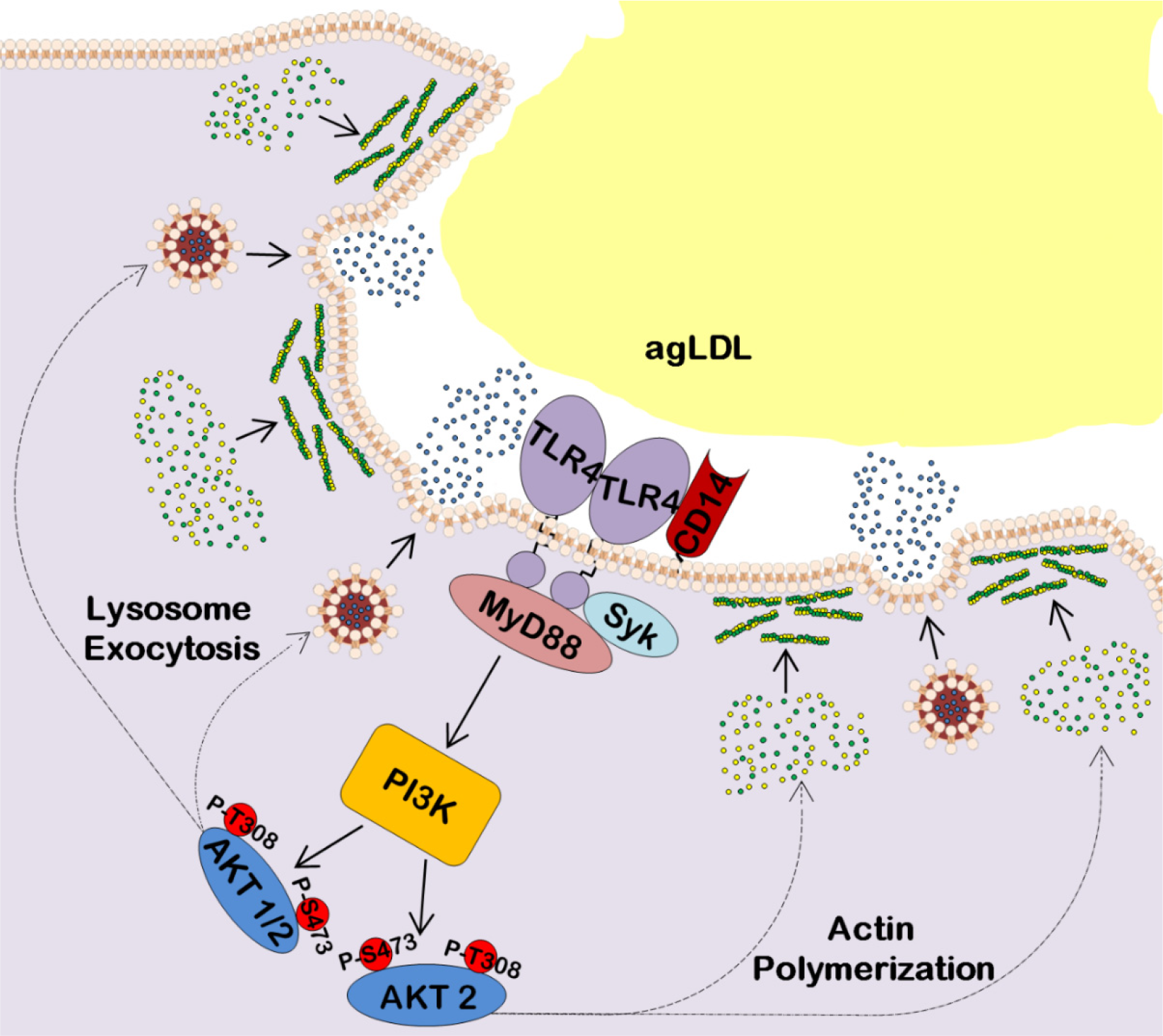
Graphic Abstract. Schematic of a signaling cascade involving TLR4, CD14, Syk, MyD88, PI3K, and AKT1/2 that is activated upon macrophage interaction with agLDL. AgLDL induced TLR4 activation triggers LS formation, along with MyD88, CD14 and SYK. Macrophage TLR4/MyD88 signaling leads to PI3 kinase activation, phosphorylation of AKT and generation of PIP_3_. Activation of AKT2 triggers local F-actin polymerization which would lead to the formation of hydrolytic compartments around agLDL. Activation of both AKT1/2 stimulates lysosomal fusion with the plasma membrane, thereby releasing lysosomal enzymes (lysosome exocytosis). These released enzymes, such as lysosomal acid lipase, would participate in the extracellular breakdown of agLDL (exophagy). This process would lead to macrophage intake of agLDL.

