## Supplemental Figures I - VIII for "TLR4-dependent signaling drives extracellular catabolism of low-density lipoprotein aggregates"

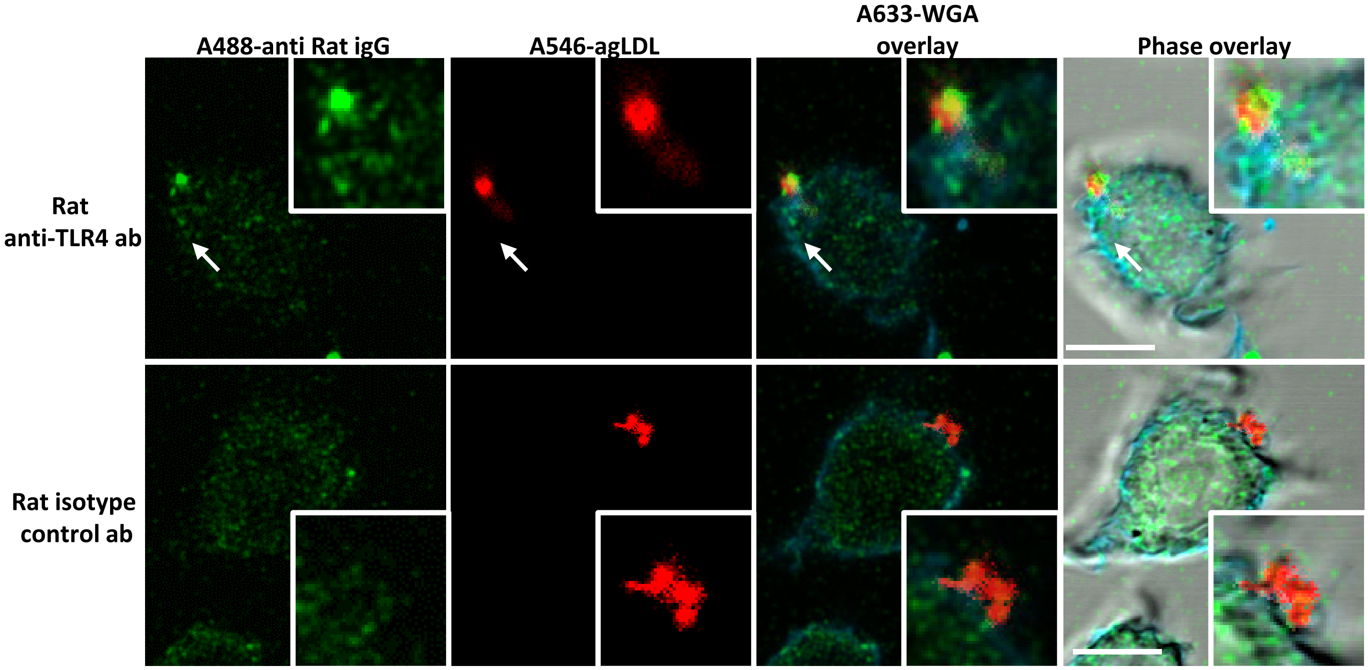


**Figure I. TLR4 clusters in the vicinity of the LS.** RAW264.7 macrophages were treated with A546-AgLDL for 1 hr and fixed using 3% PFA/0.5% glutaraldehyde. TLR4 was detected using a Rat anti-TLR4 antibody, followed by A488-anti Rat secondary antibody. Rat isotype control antibody was used to assess non-specific antibody binding. Cell plasma membrane was stained using A633-WGA. Arrow shows TLR4 clustering at the LS. Scale bars 20 µm.

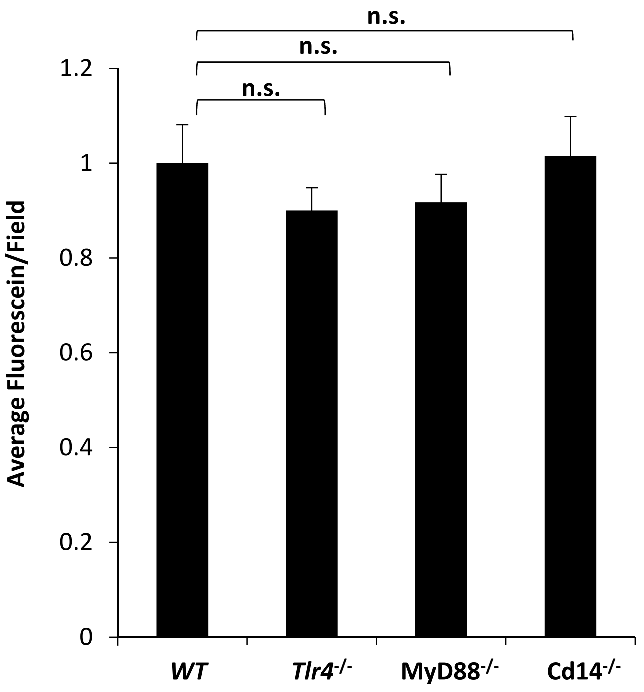


**Figure II. Biotin-fluorescein-dextran loading control.** No difference was seen in the pinocytosis of biotin-fluroescein-dextran between WT, *Tlr4*$^{-/-}$, *Myd88*$^{-/-}$ and *Cd14*$^{-/-}$ macrophages. Data were compiled from at least 3 independent experiments per condition. Error bars represent the SEM. n.s. is non significant.

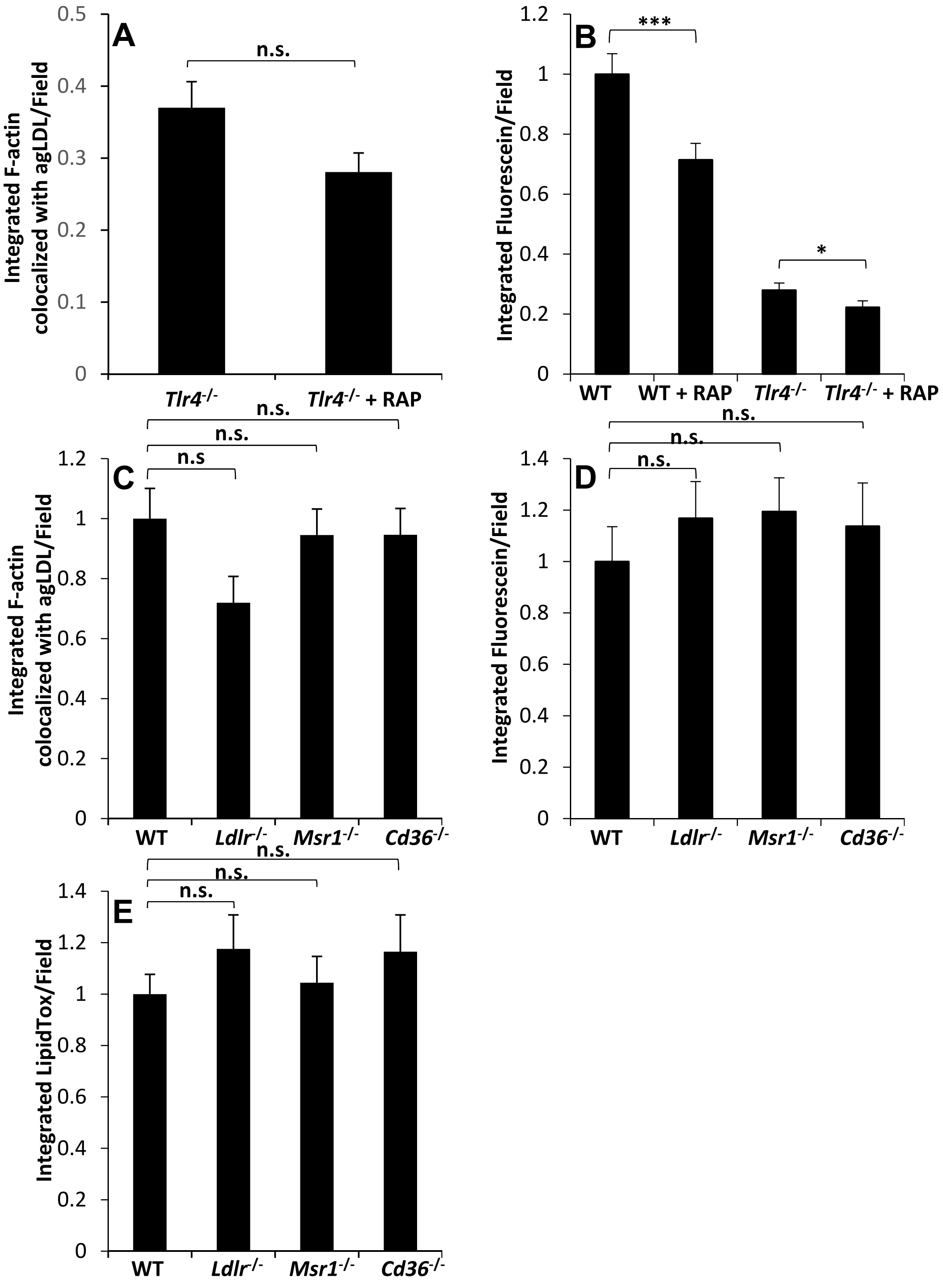


**Figure III. LRP can promote lysosome exocytosis but LDLr, MSR1 and CD36 are not involved in actin polymerization, lysosome exocytosis and foam cell formation in response to agLDL.** (A) *Tlr4*$^{-/-}$ BMMs were left untreated or pre-treated with RAP for 1 hr prior to incubation with A546-agLDL for 1 hr in the presence of RAP. Local F-actin rich structures used to form the compartment were quantified via confocal microscopy of A488-phalloidin stained samples and normalized to WT BMMs in the same experiments. (B) WT and *Tlr4*$^{-/-}$ BMMs with their lysosomes loaded with biotin-fluorescein-dextran were left untreated or pre-treated with RAP for 1 hr prior to incubation with streptavidin-A546-agLDL for 90 min in the presence of RAP. Exocytosis of biotin-fluorescein-dextran to aggregate containing compartments was quantified via confocal microscopy. (C) Quantification of the local F-actin rich structures used to form the lysosomal synapse in WT, *Ldlr*$^{-/-}$, *Msr1*$^{-/-}$ and *Cd36*$^{-/-}$ BMMs. (D) Quantification of lysosome exocytosis to aggregate containing compartments in WT, *Ldlr*$^{-/-}$, *Msr1*$^{-/-}$ and *Cd36*$^{-/-}$ BMMs. (E) Quantification of foam cell formation in WT, *Ldlr*$^{-/-}$, *Msr1*$^{-/-}$ and *Cd36*$^{-/-}$ BMMs incubated with A546-agLDL for 12 hrs, fixed and stained with LipidTOX green. Data were compiled from at least 3 independent experiments per condition. Error bars represent the SEM. * $p \leq 0.05$, *** $p \leq 0.001$. N.s. Not statistically significant.

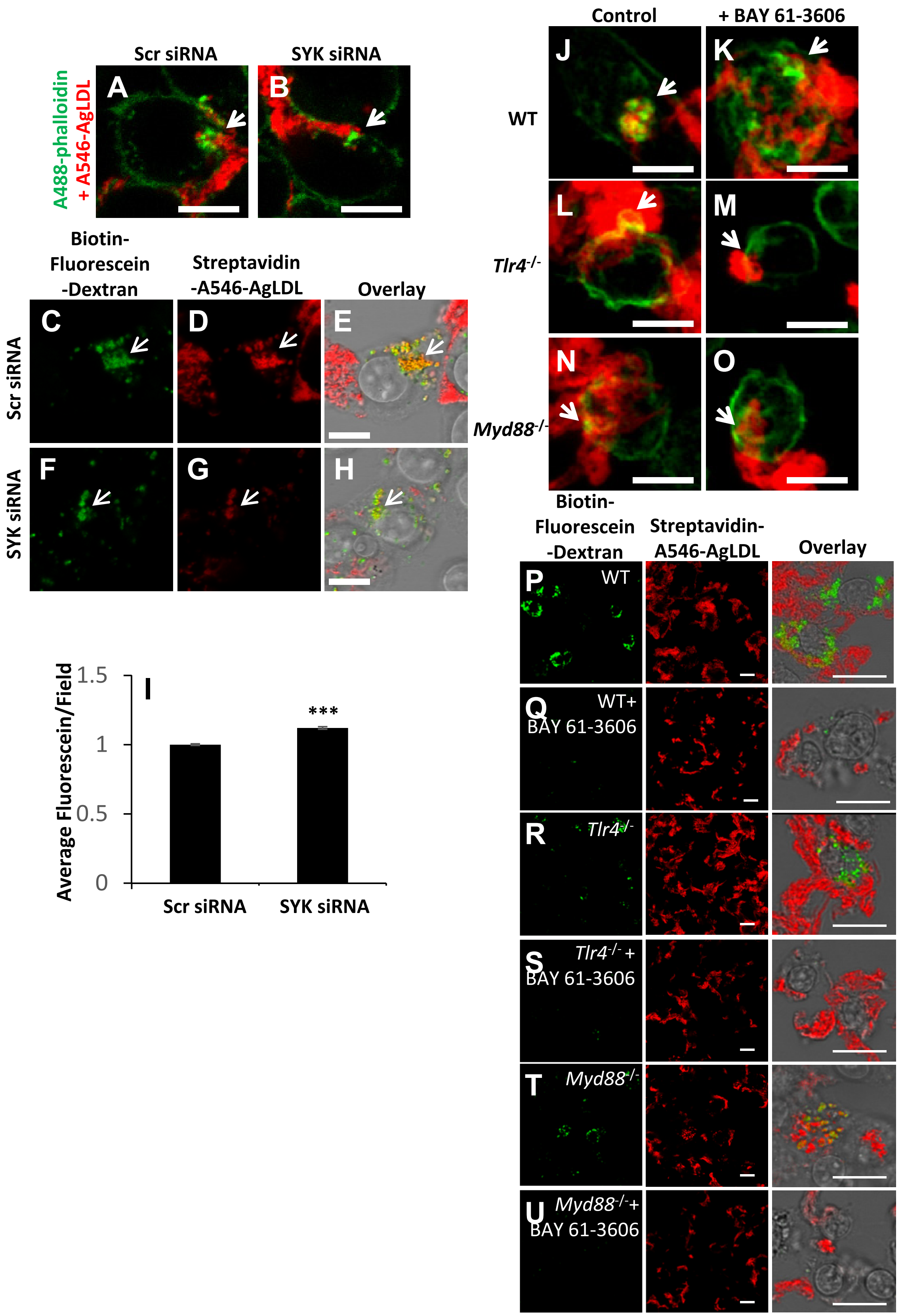


**Figure IV. SYK can promote actin polymerization and lysosome exocytosis in response to agLDL.** (A-B) Representative images from Figure 3C showing actin polymerization at the LS in Scr siRNA or SYK siRNA transfected J774 cells. (C-H) Representative images from Figure 3D showing lysosome exocytosis at the LS in Scr siRNA or SYK siRNA transfected J774 cells. (I) Scr siRNA or SYK siRNA transfected J774 cells were treated overnight with fluorescein-biotin-dextran and dextran loading in lysosomes was quantified via confocal microscopy. (J-O) Representative images from Figure 3E showing actin polymerization at the LS in WT, *Tlr4-/-* or *MyD88-/-* BMMs with or without BAY 61-3606 treatment to inhibit SYK. (P-U) Representative images from Figure 3F showing lysosome exocytosis at the LS in WT, *Tlr4-/-* or *MyD88-/-* BMMs with or without BAY 61-3606 treatment to inhibit SYK. Scale bars 20 μm. Data were compiled from 3 independent experiments. Error bars represent the SEM. *** p ≤ 0.001.

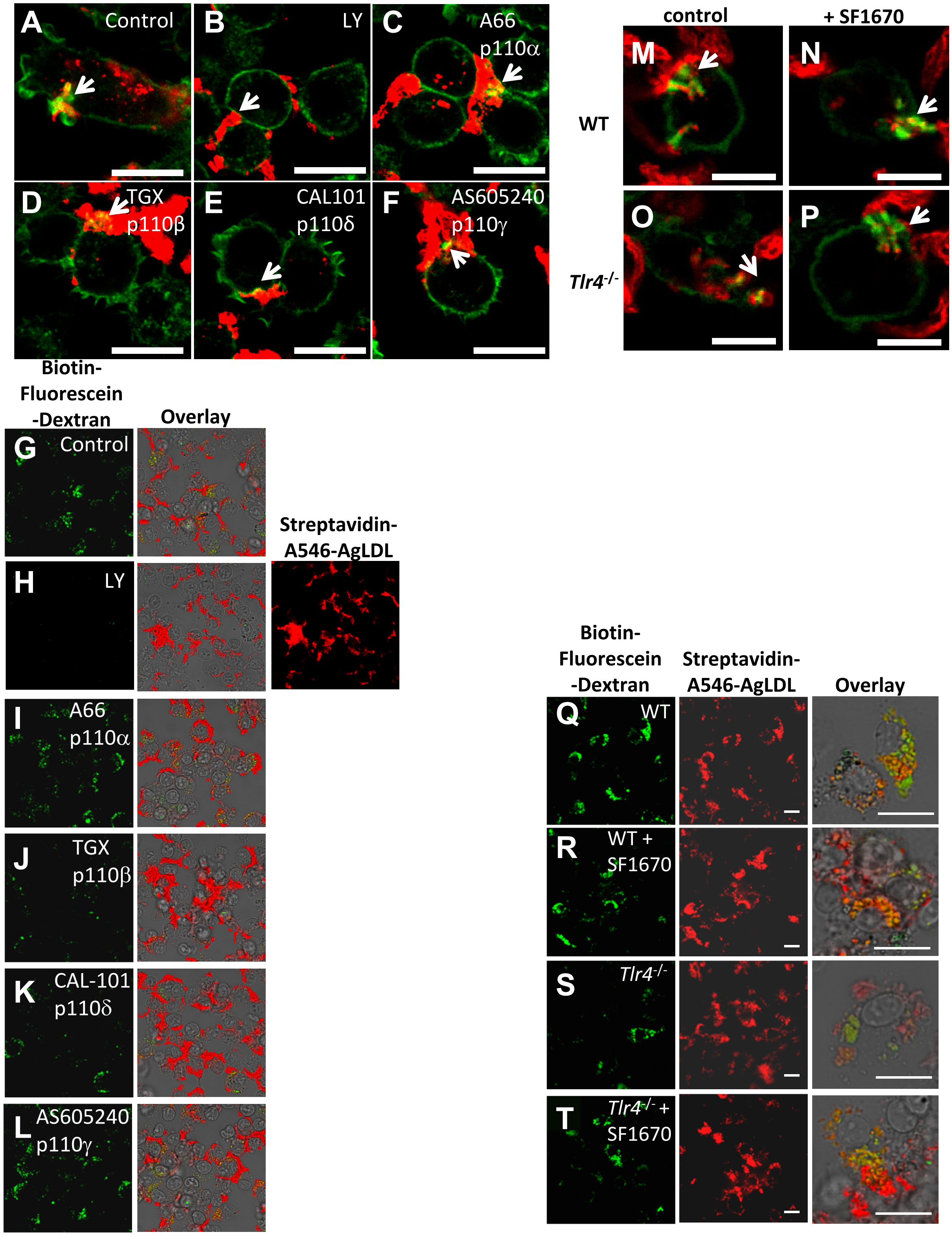


**Figure V. PI3 kinase regulates actin polymerization and lysosome exocytosis to the LS.** (A-F) Representative images from Figure 4A showing actin polymerization at the LS in J774 cells pre-treated with DMSO (control), LY294002 (50 μM), p110α inhibitor A66 (8 μM), p110β inhibitor TGX-221 (2 μM), p110δ inhibitor CAL-101 (2 μM) and p110γ inhibitor AS-605240 (2 μM) for 1 hr prior to treatment with A546-agLDL for 1 hr in the presence of inhibitors. (G-L) Representative images from Figure 4B showing lysosome exocytosis at the LS in J774 cells pre-treated with DMSO (control), LY294002 (50 μM), p110α inhibitor A66 (8 μM), p110β inhibitor TGX-221 (2 μM), p110δ inhibitor CAL-101 (2 μM) and p110γ inhibitor AS-605240 (2 μM) for 1 hr prior to treatment with Streptavidin-A546-agLDL for 90 min in the presence of inhibitors. (M-P) Representative images from Figure 4E showing actin polymerization in WT and *Tlr4*$^{-/-}$ BMMs pre-treated with DMSO control or PTEN inhibitor SF1670 (1 μM) for 1 hr prior to treatment with A546-agLDL for 1 hr in the presence of inhibitors. (Q-T) Representative images from Figure 4F showing WT and *Tlr4*$^{-/-}$ BMMs pre-treated with DMSO control or PTEN inhibitor SF1670 (1 μM) for 1 hr prior to treatment with Streptavidin-A546-agLDL for 90 min in the presence of inhibitors. Scale bars 20 μm.

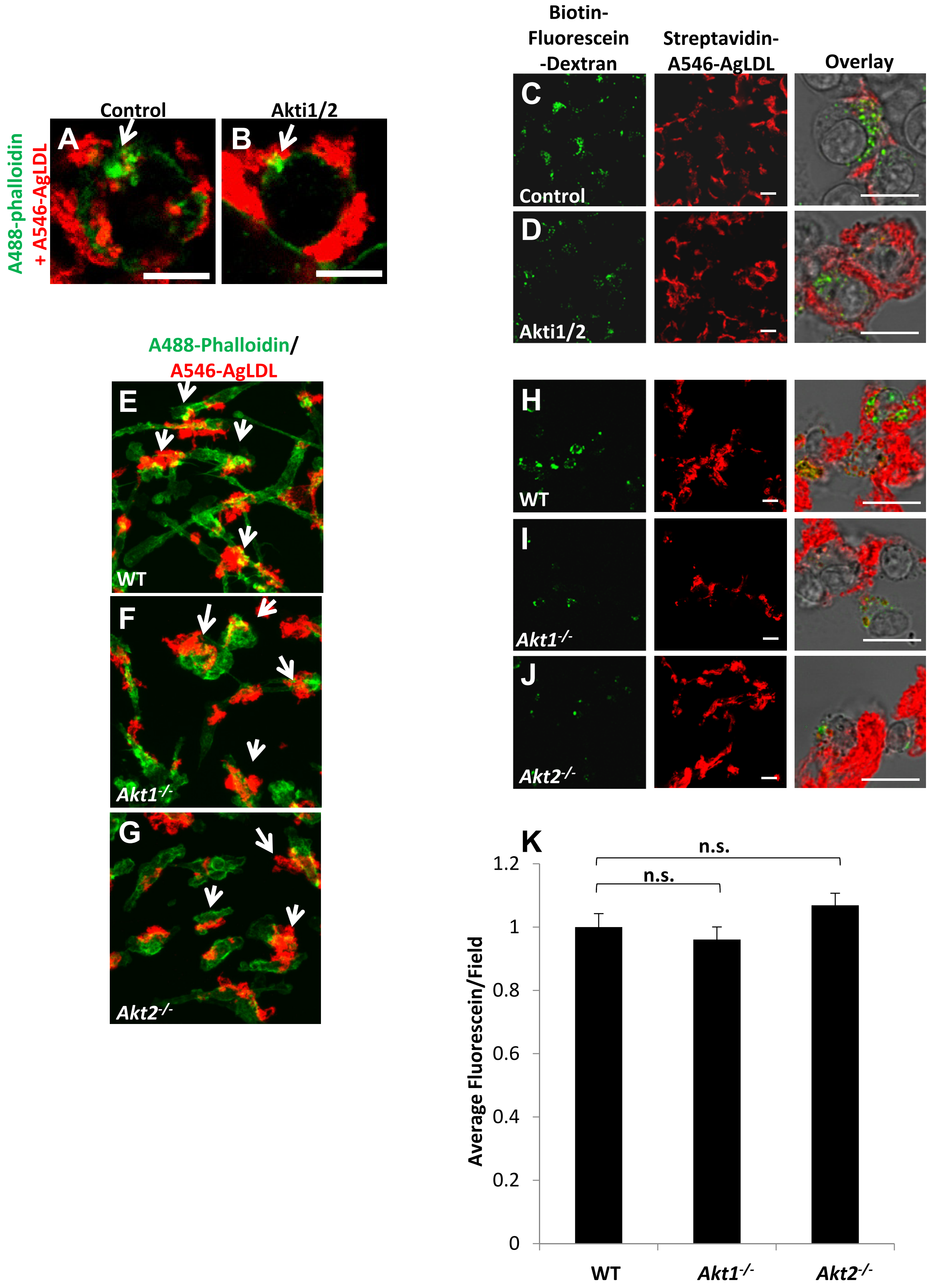


**Figure VI. Akt can promote actin polymerization and lysosome exocytosis in response to agLDL.** (A-B) Representative images from Figure 5A showing actin polymerization at the LS in J774 cells pre-treated with DMSO (control) or Akti1/2 for 1 hr prior to treatment with A546-agLDL for 1 hr in the presence of inhibitor. (C-D) Representative images from Figure 5B showing lysosome exocytosis at the LS in J774 cells pre-treated with DMSO (control) or Akti1/2 for 1 hr prior to treatment with Streptavidin-A546-agLDL for 90 min in the presence of inhibitor. (E-G) Representative images from Figure 5C showing actin polymerization at the LS in WT, *Akt1*$^{-/-}$ and *Akt2*$^{-/-}$ BMM treated with A546-agLDL for 1 hr. (H-J) Representative images from Figure 5D showing lysosome exocytosis at the LS in WT, Akt1$^{-/-}$ and Akt2$^{-/-}$ BMM treated with Streptavidin-A546-agLDL for 90 min. (K) WT, *Akt1*$^{-/-}$and *Akt2*$^{-/-}$ BMMs were treated overnight with fluorescein-biotin-dextran and dextran loading in lysosomes was quantified via confocal microscopy Scale bars 20 µm. n.s. is non significant.

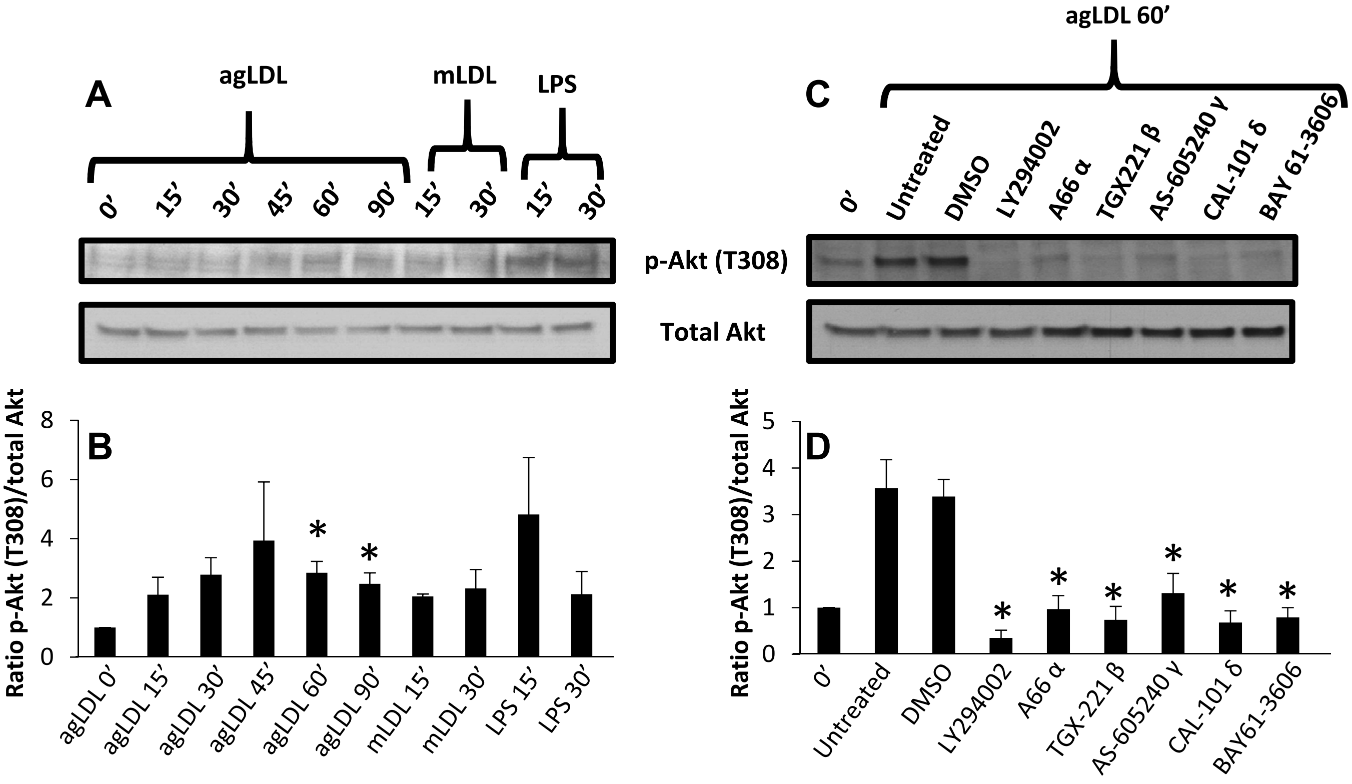


**Figure VII. AgLDL-stimulated Akt activation occurs in a PI3K/SYK dependent manner.** (A) J774 cells were treated with agLDL (1 mg/mL), mLDL (1 mg/mL) or LPS (100 ng/mL) for indicated periods of time. Ratios of phosphorylated to total protein were obtained by densitometry analysis for (B) Akt (T308). (C) J774 cells were pre-treated for 1 hr with various inhibitors (LY294002 50 μM, A66 8 μM, TGX-221 2 μM, AS-605240 2 μM, CAL-101 2 μM, and BAY 61-3606 5 μM), left untreated or treated with DMSO control, and subsequently incubated with agLDL (1 mg/mL) for 1 hr in the presence of inhibitors. Ratios of phosphorylated to total protein were obtained by densitometry analysis for (D) Akt (T308). Data were compiled from 3 independent experiments. Error bars represent the SEM. * $p < 0.05$.

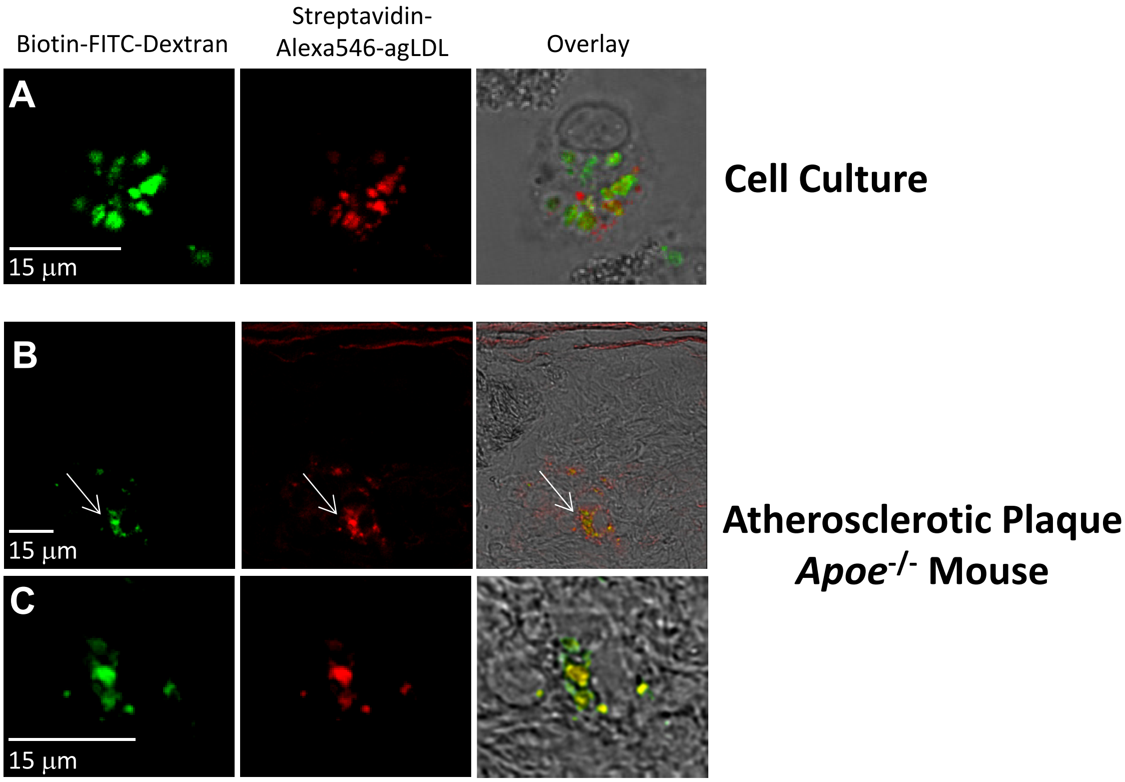


**Figure VIII. Comparison of LS morphology in vitro and in vivo.** (A) J774 cells with their lysosomes loaded with biotin-fluorescein-dextran were incubated with streptavidin-A546-agLDL for 90 min, fixation and permeabilized. (B, C) Apoe$^{-/-}$ mice on a HFD were injected with streptavidin-A546-LDL 1 days prior to injection with BMMs with their lysosomes loaded with biotin-fluorescein-dextran. 3 days after adoptive transfer mice were sacrificed, and the aortas were harvested, sectioned and examined by confocal microscopy.
